# The human-restricted oncoprotein POU5F1B enhances cell invasiveness through plasma membrane remodeling

**DOI:** 10.1101/2023.10.30.564723

**Authors:** Sandra Offner, Laurence Abrami, Elisa Burri, Adrien Mery, Barney Drake, Jialin Shi, Eduard M Unterauer, Julien Duc, Evarist Planet, Ralf Jungmann, Alexandre Reymond, Georg Fantner, Mahmut Selman Sakar, Bernhard Wehrle-Haller, F. Gisou van der Goot, Didier Trono, Laia Simó-Riudalbas

**Author notes:** **Declaration of interests:** pending patent application -POU5F1B inhibitors-. Inventors: Laia Simó-Riudalbas, Didier Trono, Gisou van der Goot, Laurence Abrami, Sandra Offner.

## Abstract

Evolution confers new species with distinctive biological features that can translate in either purely mechanistic speciation or novel phenotypes. Retrotransposition in the last Hominidae common ancestor of the pluripotency regulator *POU5F1/OCT4* led to human POU5F1B, which promotes gastrointestinal cancer growth and metastasis through unknown mechanisms. Here, we show that POU5F1B fosters cell invasiveness by inducing plasma membrane remodeling. This ability exquisitely depends on a series of post-translational modifications. Ubiquitination of two lysine residues found exclusively in human POU5F1B results in its cytoplasmic retention, contrasting with the nuclear POU5F1. ZDHHC17-mediated S-acylation then triggers POU5F1B association with membrane nanodomains enriched in cell adhesion and signaling molecules. This is accompanied by the cell surface clustering and accelerated turnover of focal adhesion proteins, with enhanced cell invasiveness. Finally, screening for inducers of POU5F1B degradation, we found its stability to depend critically on Rho-associated protein kinases, revealing potential avenues for the treatment of POU5F1B-expressing tumors.

## Introduction

Gene duplication is a prominent mechanism of emergence of new gene functions^1^. Transposable elements (TEs) contribute to this process through the generation of mRNA-derived gene duplicates, known as retrogenes. This usually occurs when the reverse transcription machinery of a TE, typically a LINE-1 retrotransposon, captures a cellular mRNA to yield a DNA copy that is then integrated elsewhere in the genome. Long collectively dismissed as non-functional “processed pseudogenes”, retrocopies are increasingly recognized as having contributed a large number of bona fide retrogenes notably in mammals and fruit flies^2–6^. The human genome contains several thousand readily identifiable retrocopies, a few hundreds of which are robustly expressed^7,8^. In most cases, these encode proteins identical to the product of the parental gene, albeit along distinct expression patterns since retrocopies are generally devoid of the source gene regulatory sequences. In a few cases, though, retrogenes undergo neofunctionalization. A typical example is the accumulation in elephant of 19 retrocopies of the tumor-suppressor gene *TP53*, some of which encode gain-of-function proteins responsible for enhanced apoptotic response to DNA damage^9–11^. Both the increased levels of TP53 and the presence of these super-protective variants likely contribute to the rarity of cancer in this species.

The gene encoding OCT4 (octamer-binding protein 4), the well characterized homeodomain transcription factor of the POU (Pit-Oct-Unc) family involved in the self-renewal of embryonic stem cells and in early embryonic development, underwent a series of retroposition events in primates. This resulted in the presence of 6 *POU5F1/OCT4* retrocopies in the human genome, in addition to the parental gene. The youngest, *POU5F1B*, emerged in the last common ancestor of great apes, some 12 million years ago, and is located on chromosome 8q24, proximal to the *MYC* locus. It is transcribed by activation of upstream transposable element-embedded regulatory sequences in several human solid tumors including colon, stomach, breast, prostate and uterus^12^. Its expression is a negative prognostic factor in colorectal^12^, gastric^13^ and hepatocellular^14^ malignancies and is associated with advanced histological grades of cervical cancer^15^, where the *POU5F1B* locus is a papillomavirus integration hotspot^16^. Supporting its role as a cancer-promoting factor, the POU5F1B protein stimulates the proliferation of colorectal^12^, gastric^13^ and hepatocellular^14^ carcinoma cells *in vitro* and increases the growth^12,13^ and metastatic potential^12^ of human gastrointestinal tumor cells in mouse xenotransplantation experiments.

It has been assumed that POU5F1B, like its parental gene OCT4, acts as a stemness-promoting transcription factor. However, we uncovered that despite minor sequence differences, it does not localize to the nucleus but rather to cellular membranes and associates with protein kinases and cytoskeleton-related molecules^12^. We also found that POU5F1B induces intracellular signaling events and the release of trans-acting factors involved in cell growth and cell adhesion^12^. The present study was undertaken to explore further the mechanism of POU5F1B oncogenic action, and notably how it promotes metastases. It reveals that POU5F1B oncogenic function depends on sequential post-translational modifications, some targeting residues unique to the human protein, and on the induction of protein-enriched membrane nanodomains, with augmented cell invasiveness. We also discovered that POU5F1B-induced malignant phenotypes can be reverted by blocking ROCK protein kinases. Thus, POU5F1B is a human-unique oncoprotein that acts by unprecedented mechanisms and can be inhibited through pharmacological approaches.

## Results

### POU5F1B cytoplasmic localization and function depend on ubiquitination of human-restricted lysine residues

POU5F1/OCT4 and POU5F1B differ by only 15 residues out of 359/360, 5 of which are unique to human POU5F1B, that is, absent from the protein expressed by other great apes (Fig. 1A, Supp. Fig. 1). Yet POU5F1/OCT4 localizes to the nucleus, while POU5F1B accumulates in the cytoplasm^12^. We sought to explain this difference by expressing wild-type and mutated forms of the oncoprotein in SW480 colorectal cancer (CRC) cells. Amongst the 5 human-specific residues of POU5F1B, lysine K135 and K182 stood out as possible targets of ubiquitination, a post-translational modification that can influence the stability, activity and subcellular localization of proteins^17^ (Fig. 1A, Supp. Fig. 1). Confirming this hypothesis, we determined that POU5F1B is ubiquitinated and that this modification is largely suppressed when K135 and K182 were changed to glutamic acid and threonine, respectively, as found in POU5F1, or when K182 was substituted by an alanine (Fig. 1B, 1C). Remarkably, the single and double (K135E/K182T) lysine mutants localized to the nucleus (Fig. 1D, 1E). Consistently, inhibiting the ubiquitin activating enzyme with TAK-243^18^ triggered the nuclear localization of wild type POU5F1B (Fig. 1D, 1E). We next tested the importance of the cytoplasmic localization of POU5F1B for its ability to promote growth. Replacing either of the two lysine residues specific of POU5F1B by the corresponding amino acid of POU5F1/OCT4 abrogated its growth-promoting action on CRC cells (Fig. 1F).

**Figure 1.**
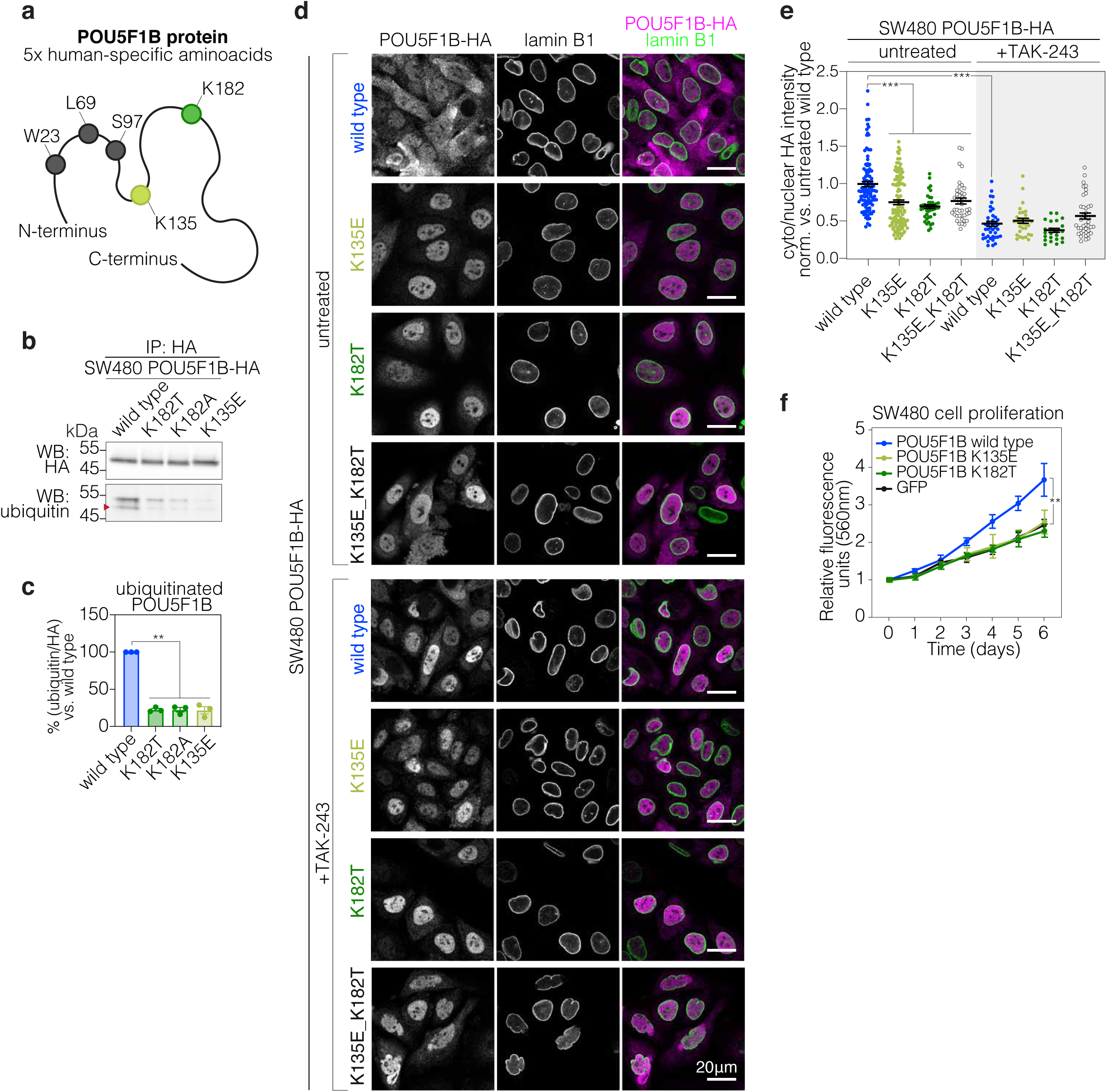
POU5F1B cytoplasmic localization and growth-stimulating effects depend on ubiquitination of human-restricted lysine residues. **a,** Schematic representation of human POU5F1B (359 aa) protein, with the five human-specific amino acids highlighted. K135 and K182 in green are focus of our study. **b,** Immunoprecipitation (IP) and western blot (WB) analysis of HA-tagged wild type (WT)-, K182T-, K182A-, and K135E-POU5F1B in SW480 cells. **c,** Quantification of western blot bands shown in (b) (wild type vs. K182T P=5.1e-04; vs. K182A P=1.6e-03; vs. K135E P=3.9e-03 by t-test). **d,** Representative immunofluorescence-confocal microscopy of untreated and TAK-243 (1uM, 5h) treated POU5F1B-HA wild type-, K135E-, K182T-, K135E-K182T-overexpressing SW480 cells, with HA in magenta and the nuclear membrane marker lamin-B1 in green. **e,** Ratio cytoplasm/nucelus of HA signal intensity from cells presented in (d). Each data point was normalized to the mean intensity of untreated POU5F1B-HA wild type cells (untreated wild type vs. K135E P=4.4e-07; vs. K182T P=2.43e-08; vs. K135E_K182T P=6.42e-05; vs. TAK243 wild type P=1.12e-15 by Wilcoxon test). **f,** PrestoBlue cell proliferation assay in POU5F1B-wild type-, -K135E-, -K182T-, and GFP-overexpressing SW480 cells. (n = 3 independent experiments, P = 0.001 by Wilcoxon test). ** pvalue<0.01, *** pvalue<0.001.

### ZDHHC17-mediated acylation is necessary for POU5F1B action on membrane nanodomains

We previously reported POU5F1B enrichment in detergent resistant membrane (DRM) fractions of CRC cells^12^. Membrane association of soluble proteins can be induced by covalent linkage of fatty acids, amongst which S-acylation (commonly referred to as S-palmitoylation) plays a particular role in recruitment to DRM subdomains^19,20^. Using the acyl-resin assisted capture (Acylrac) method, whereby S-acylated proteins are trapped on thiol-reactive sepharose beads, we could demonstrate that POU5F1B is S-acylated, and that this modification is reduced by the K135E and K182T mutations (Fig. 2A, 2B). We also found that these amino acid substitutions accordingly impaired POU5F1B association to DRM, assessed using the gradient-based DRM subcellular fractionation as a biochemical readout of cholesterol- and sphingolipid-rich nanodomains^21,22^ (Fig. 2C, 2D). Moreover, we observed that wild type POU5F1B expression triggered a significant (>20%) redistribution of transferrin receptor (TFRC), normally a marker of detergent soluble membranes, into DRMs^12^ (Fig. 2C, 2D). These findings suggest that POU5F1B induces the formation of non-canonical nanodomains in CRC cells. To explore further this phenomenon, we isolated DRMs from SW480 cells overexpressing wild type POU5F1B, its K135E mutant or GFP and analyzed the various fractions of the gradient by Coomassie staining (Supp. Fig. 2A, 2B). Expression of wild type POU5F1B markedly increased the protein content of DRMs, when compared to cells expressing either the K135E mutant or GFP (Supp. Fig. 2A, 2B).

**Figure 2.**
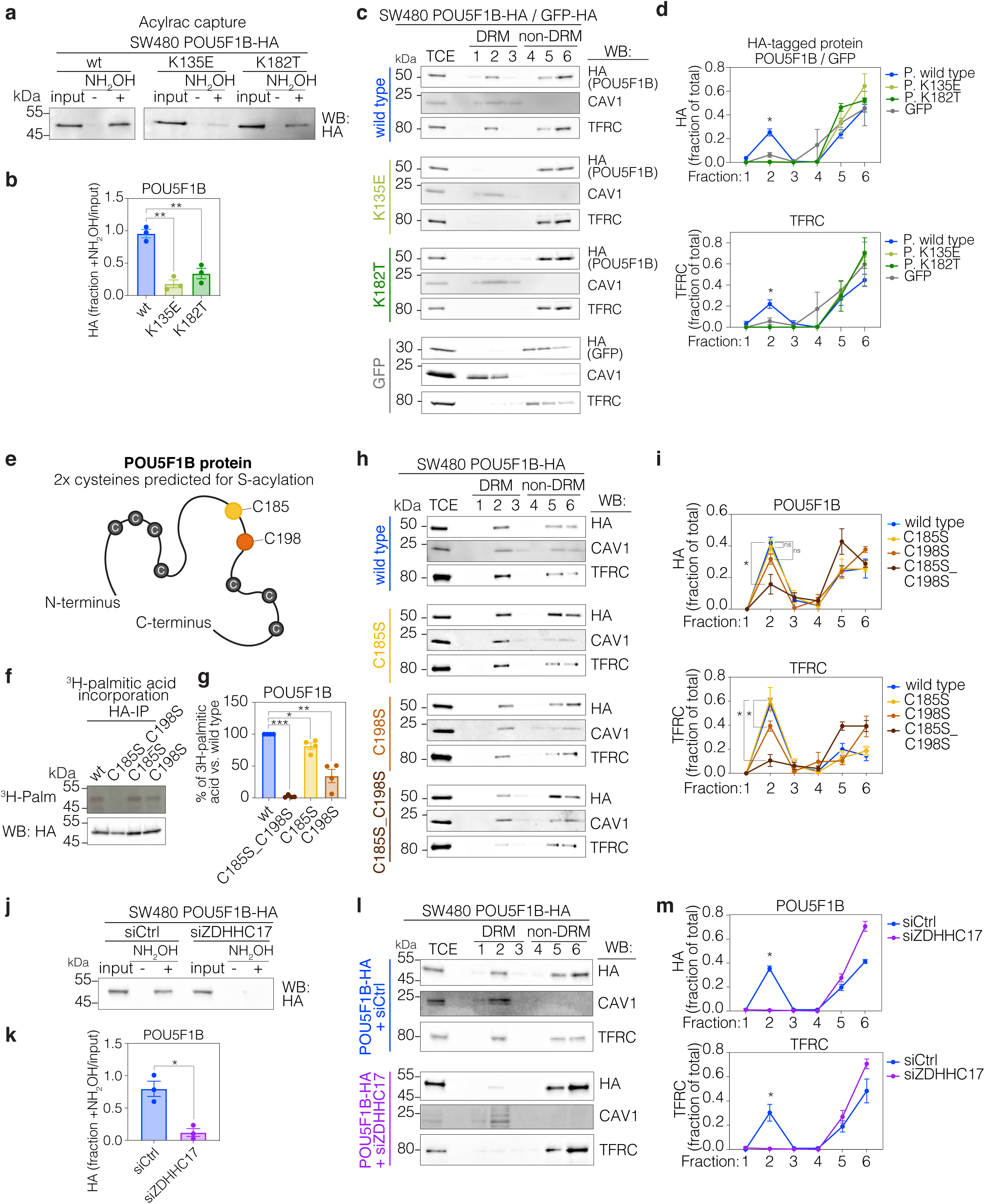
ZDHHC17-mediated acylation is necessary for POU5F1B action on membrane nanodomains. **a,** Acylrac capture assay in POU5F1B-HA wild type-, K135E-, K182T-overexpressing SW480 cells. Western blot (WB) anti-HA of total cell extracts (input), S-acylated (plus hydroxylamine +NH_2_OH) and control fractions (–NH_2_OH). **b,** Ratio of HA signal in S-acylated fraction/total cell extracts from three independent acylrac experiments (wild type vs. K135E P=0.001; vs. K182T P=0.004 by t-test). **c,** Isolation of detergent-resistant membranes (DRMs) from POU5F1B-HA-wild type-, -K135E-, -K182T-, and GFP-overexpressing SW480 cells. 1-3 correspond to insoluble fractions, 4-6 to soluble fractions, DRMs being traditionally found in fraction 2. Caveolin1 (CAV1) and transferrin receptor (TFRC) are used as controls for insoluble and soluble fractions, respectively (representative blot out of three independent experiments). **d,** Quantification of western blot in (c) normalized within each experiment (Fraction 2 in HA blot: wild type vs. K135E P=0.013; vs. K182T P=0.012; vs. GFP P=0.009; fraction 2 in TFRC blot: wild type vs. K135E P=0.035; vs. K182T P=0.030; vs. GFP P=0.040 by t-test). **e,** Schematic representation of human POU5F1B protein, with the nine cysteines highlighted. C198 is predicted for S-aclyation by SwissPalm and C185 by physical proximity to C198. **f,** Autoradiography (^3^H-Palm) and WB of immunoprecitated HA-tagged wild type (wt)-, C185S_C198S-, C185S-, C198S-POU5F1B in ^3^H-palmitic acid-labeled SW480 cells. **g,** Quantification of bands in (f) (n=3 independent experiments; P=2.38e-06, P=0.034, P=0.008, respectively, by t-test). **h,** Isolation of DRMs from cells in (f). **i,** Quantification of western blot bands in (h) (Fraction 2 in POU5F1B blot: POU5F1B wild type vs. C185S P=0.65 ; vs. C198S P=0.079 ; vs. C185S_C198S P=0.015; fraction 2 in TFRC blot: POU5F1B wild type vs. C185S P=0.765 ; vs. C198S P=0.048 ; vs. C185S_C198S P=0.025 by t-test). **j,** Acylrac capture assay in siCtrl-, and siZDHHC17-POU5F1B-HA SW480 cells. **k,** Quantification of blot in (j) (P=0.014 by t-test). **l,** Isolation of DRMs from cells in (j). **m,** Quantification of western blot bands in (l) (Fraction 2 in POU5F1B blot P=0.002; fraction 2 in TFRC blot P=0.049 by t-test). ns, not significant, * pvalue<0.05, ** pvalue<0.01, *** pvalue<0.001

S-acylation occurs on cysteines. According to the SwissPalm database (https://swisspalm.org/proteins/Q06416), out of nine such residues present in POU5F1B, only cysteine198 is predicted to be acylated with a high confidence score. We mutated to serine Cys198, as well as Cys185 given its relative proximity (Fig. 2E). We evaluated the dynamics of POU5F1B S-acylation by monitoring the incorporation of radioactive 3H-palmitic acid using immunoprecipitation and autoradiographic analysis. The results confirmed POU5F1B S-acylation (Fig. 2F, 2G), and revealed that this modification was affected more importantly by the C198S than the C185S mutation, yet was dramatically reduced in the C185S/C198S double mutant, suggesting cooperative S-acylation. Correspondingly, this double mutant only marginally associated with DRMs where it failed to induce TFRC recruitment (Fig. 2H, 2I) and protein accumulation (Supp. Fig. 2C, 2D).

Protein S-acylation is mediated by protein S-acyltransferases characterized by an aspartate-histidine-histidine-cysteine (ZDHHC) domain. The human genome encodes 23 ZDHHCs, the substrate specificity of which is partly dictated by their subcellular localization^23^. Using siRNAs against each one of these 23 enzymes alone or in combinations, we identified ZDHHC17 as responsible for POU5F1B S-acylation (Fig. 2J, 2K; Supp. Fig. 2E). We next tested whether zDHHC17-mediated acylation affects the membrane association of POU5F1B. We found the DRM-association of POU5F1B to be drastically reduced upon zDHHC17 silencing, which also prevented TFRC redistribution (Fig. 2L, 2M) and protein accumulation (Supp. Fig. 2F, 2G) in DRMs of POU5F1B-expressing cells. In contrast, depleting the acyltransferase had no effect on DRMs in control cells (Supp. Fig. 2F, 2G.

Altogether, these observations indicated that POU5F1B associates with cholesterol- and sphingolipid-enriched detergent-resistant nanodomains of the plasma membrane through ZDHHC17-mediated S-acylation, and that it increases their protein recruiting capacity even extending it to proteins such as the transferrin receptor, normally excluded from these domains.

### POU5F1B triggers the redistribution of cell adhesion proteins and changes in cell morphology

The POU5F1B-induced modifications of cell membranes suggested that it might also endow the cell surface with special topographical features. To test this hypothesis, we applied scanning ion conductance microscopy (SICM)^24^ (Fig. 3A) to measure cell height and roughness in living GFP-, POU5F1B-, and K135E-POU5F1B-expressing SW480 cells (Fig. 3B). The results indicated that wild type POU5F1B-expressing cells were taller than the GFP control and had a rougher surface over the cell body (Fig 3C, 3D) and borders (Fig. 3E) than the two other cell types tested. Images further revealed the presence of membrane protrusions projected upward in the z axis in POU5F1B-expressing cells (Fig. 3B), consistent with our previous finding that levels of ruffle-localized proteins such as SLC9A3R1, CD2AP, RDK and EZR are increased in POU5F1B-expressing cells^12^.

**Figure 3.**
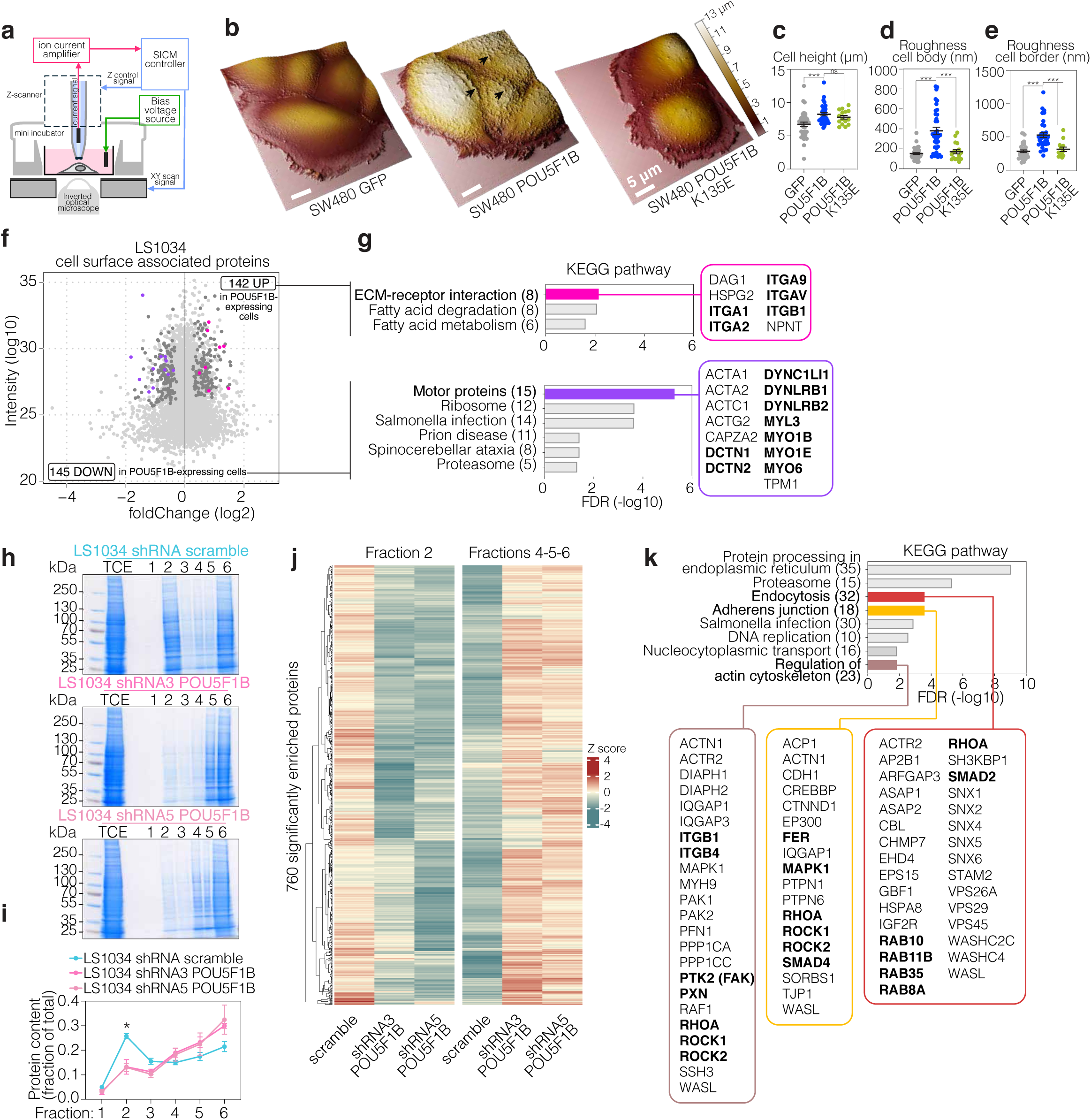
POU5F1B triggers the redistribution of cell adhesion proteins and changes in cell morphology. **a,** Scheme ilustrating the scanning ion conductance microscopy (SICM) tehcnique to study the surface morphology of cells. **b,** Representative SICM 3D images showing cell topography from GFP-, POU5F1B- and POU5F1B-K135E-expressing SW480 cells. Color scale from 0 to 13 µm reveals cell heigth values. Black arrows indicate protrusions projected upward in the z axis. **c,** Cell height (µm) (POU5F1B vs. GFP P=0.0005; vs. K135E P=0.35 by Wilcoxon test), **d,** roughness of cell body (nm) (POU5F1B vs. GFP P=1.42e-07; vs. K135E P=0.0002 by Wilcoxon test), **e,** and roughness of cell border (nm) (POU5F1B vs. GFP P=3.63e-07; vs. K135E P=0.0001 by Wilcoxon test) from cells in b. **f,** MA plot depicting relative abundance and average intensity of individual proteins identified by mass spectrometry from cell surface-associated proteins of sh-scramble vs. shRNA3 & shRNA5 LS1034 cells. Measurements were performed in independent triplicates, each dot represents a detected protein, with significantly changed ones (P < 0.05, outlier detection test as computed by MaxQuant) in dark grey and their numbers indicated in upper corners. **g,** Significantly enriched proteins were clustered in KEGG annotations using the Functional Annotation Chart from DAVID bioinformatics resources 6.8. In bold, proteins mentioned in the text. **h,** Isolation of detergent-resistant membranes (DRMs) from sh-scramble, shRNA3 & shRNA5 LS1034 cells, fractions were run on SDS-PAGE and gels were stained with Coomassie. **i,** Quantification of coomassie staining in (c) (Fraction 2 sh-scramble vs. shRNA3 P=0.018; vs. shRNA5 P=0.015 by t-test). **j,** Heatmap of mass spectrometry spectral counts from fraction 2 (F2) and fractions 4, 5, and 6 (F456), extracted from cells in (h). **k,** Significantly enriched proteins in F2 shRNA-scramble were clustered as in (g). In bold, proteins mentioned in the text. ns not significant, * pvalue<0.05, *** pvalue<0.001.

We next tested whether the physical transformation of the cell surface correlated with a change in its protein content. For this we performed surface biotinylation coupled to precipitation with streptavidin beads on control and dox-inducibly POU5F1B-depleted LS1034 CRC cells, which express this protein at baseline and were modified by lentivector-mediated drug-controllable RNA interference. Based on stringent filtering criteria (fold-change in the same direction with two different shRNAs; high intensity candidates –over quantile 25%-; and p value<0.05), we found 142 proteins enriched and 145 depleted in the surface-associated proteome of control compared with POU5F1B-depleted LS1034 cells (Fig. 3F). In their overwhelming majority (274/287), these proteins displayed no difference in total levels, indicating that POU5F1B was acting on their intracellular distribution, not on their expression level (Supp. Fig. 3A). Subjecting the subset of enriched proteins to functional enrichment analysis with the Kyoto Encyclopedia of Genes and Genomes (KEGG) using DAVID knowledgebase v2024q2^25,26^ pointed to extracellular matrix (ECM)-receptor interaction molecules as enriched features, notably ITGB1, ITGA1, ITGA2, ITGA9, and ITGAV, which are integrins involved in cell adhesion (Fig. 3G). By contrast, cytoskeletal motor proteins important for vesicle trafficking^27^, including dynein, dynactin and myosin, stood out as factors redistributed away from the surface of POU5F1B-expressing cells (Fig. 3G).

To investigate proteins enriched in POU5F1B-associated membrane nanodomains, we subjected detergent-treated protein extracts from POU5F1B-knockdown and control LS1034 cells to gradient fractionation followed by Coomassie staining (Fig. 3H, 3I) and mass spectrometry analysis (Fig. 3J). Both Coomasie staining and hierarchical clustering of mass spectrometry spectral counts from detergent insoluble F2 and soluble F4-5-6 fractions revealed that the presence of POU5F1B is associated with a higher protein content in DRM fractions (Fig. 3H, 3I, 3J), consistent with the observations made in POU5F1B-overexpressing SW480 cells (Supp. Fig. 2A). Proteins enriched in DRMs of POU5F1B^+^ cells included molecules participating in endocytosis (e.g. RAB proteins), adherent junctions and regulation of actin cytoskeleton (Fig. 3K). As expected from the strong connection between these 3 cellular functions, some proteins were classified in two or all three of these categories, e.g. the small GTPase associated with cytoskeleton dynamics and cell migration RHOA, the protein kinases involved in actin cytoskeleton organization, stress fiber and focal adhesion (FA) formation ROCK1 and ROCK2, the stimulator of actin-nucleating activity of the Arp2/3 complex WASL, and the focal adhesion components ITGB1, ITGB4, FAK and paxillin (PXN). Signaling molecules such as the signal transduction proteins activated by transforming growth factor (TGF)-beta SMAD2 and SMAD4, the serine/threonine kinase from the MAPK/ERK cascade MAPK1, and the tyrosine kinase FER, which acts downstream of several cell surface receptors, were also enriched in DRMs of POU5F1B+ cells.

POU5F1B-triggered protein redistribution was concomitant to a drop in motor proteins association with the cell surface (Fig. 3G) and a DRM enrichment of factors involved in endocytosis, such as the early endosome marker EEA1 and the recycling endosome protein RAB11B (Fig. 3K). This was confirmed by immunofluorescence analysis against early (EEA1), recycling (RAB11B) and late (LBPA, LAMP1) endosomes: while late endosomes/lysosomes were unaffected by the expression of POU5F1B, early and recycling endosomes were more abundant (Supp. Fig. 3B, 3C). Intriguingly, early endosomes were smaller in POU5F1B expressing cells, while recycling endosomes were larger. Regardless the increase in the early-recycling endosomal pathway suggests an accelerated turnover of cell surface components. Taken together, these data indicate that, in colorectal cancer cells, POU5F1B induces changes in the cell surface proteome with alterations of its 2D distribution and of the plasma membrane architecture, as well as modifications in the endosomal recycling pathway.

### POU5F1B promotes the nano-clustering, surface redistribution and increased turnover of focal adhesion proteins

Integrins, which are major components of FAs, were amongst the most surface-enriched DRM-associated proteins in POU5F1B expressing cells (Fig. 3G, 3K). Western blot analysis of total and active form of ITGB1 in the LS1034 knockdown and SW480 overexpression systems confirmed its enrichment in DRMs of POU5F1B-expressing cells (Supp. Fig. 4A, 4B). One of the primary intracellular downstream signalling mediators of integrins is the focal adhesion kinase FAK, which was also highly enriched in DRMs of POU5F1B-expressing cells (Fig. 3K). We examined the cell surface distribution of FA sites by labelling for p-Y^397^FAK, the form of FAK that becomes phosphorylated and activated upon integrin clustering^28^. To reach high <20 nm spatial resolution, we used the localization-based super-resolution approach DNA-PAINT, which achieves fluorescence separation by the transient interaction of short DNA oligos (Fig. 4A). Quantitative analysis of 75 x 75µm2 fields of view revealed a higher percentage of p-Y^397^FAK positive sites in POU5F1B-overexpressing than GFP control SW480 cells (Fig. 4B), with increased area and number of these sites (Fig. 4C). To characterize further the difference in molecular arrangements of focal adhesion sites, we performed high resolution DNA-PAINT experiments with a localization precision of ∼3.5 nm. Using DBSCAN cluster analysis we segmented p-Y^397^FAK nanoclusters within a detection range of 20nm radius in the adhesion sites (Fig. 4D). We detected a higher number of positive nanoclusters (Fig. 4E) and higher density of molecules within each nanocluster (Fig. 4F) in POU5F1B-expressing compared to control cells.

**Figure 4.**
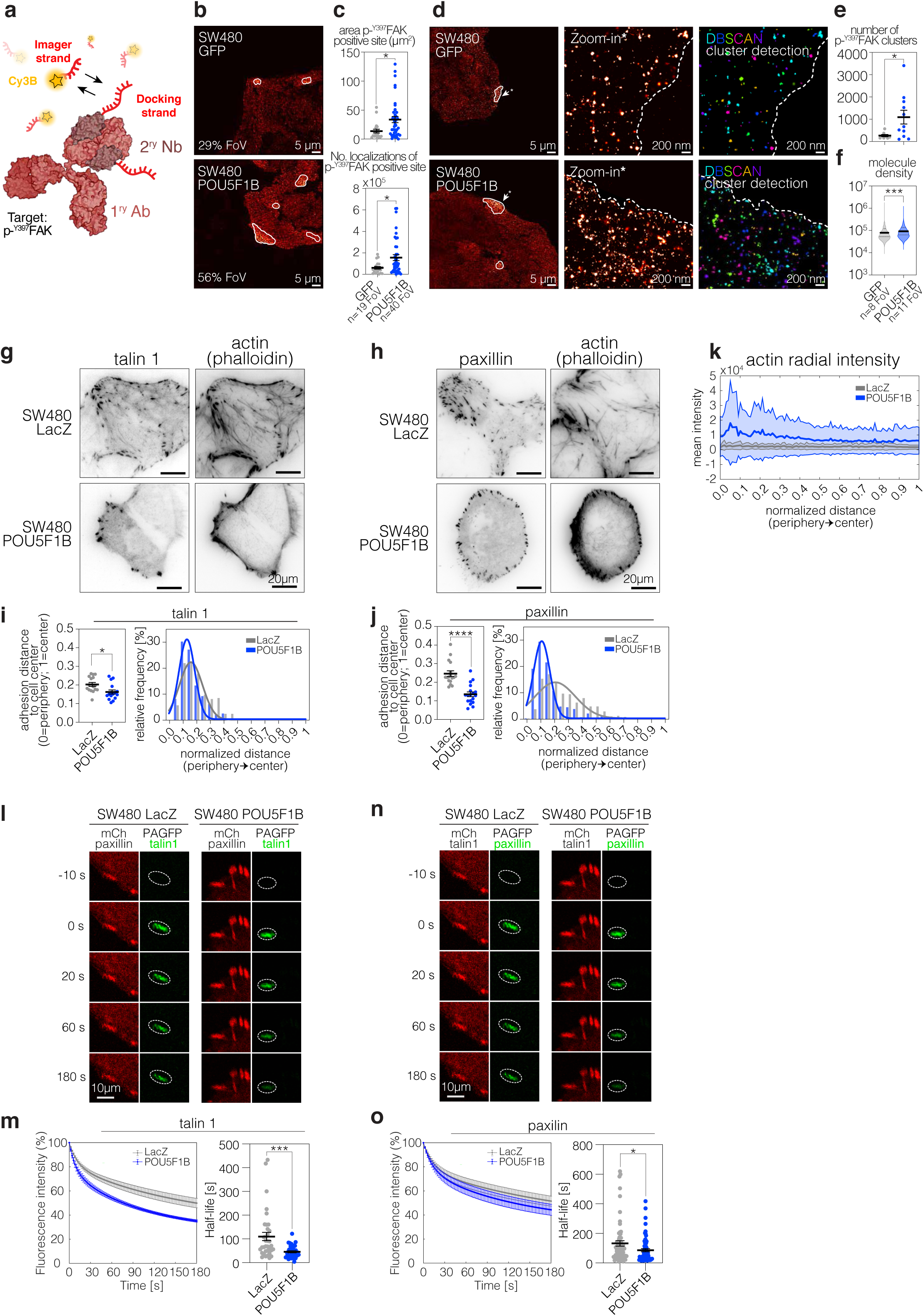
POU5F1B promotes the nano-clustering, surface redistribution and increased turnover of focal adhesion proteins. **a,** Scheme of DNA-PAINT technique targeting p-^Y397^FAK protein. Birefly, p-^Y397^FAK is labeled with a primary antibody (1ry Ab) preincubated with secondary nanobody (Nb) fused to a DNA-PAINT docking sequence (red). For readout, a complementary dye-labeled (Cy3B) imager strand is added. **b,** Representative p-^Y397^FAK-targeting DNA-PAINT images from GFPcontrol- and POU5F1B-overexpressing SW480 cells. Percentage of positive field of views (FoV) (b), area (P=0.016 by t-test) and number of localizations (P=0.026 by t-test) (**c**) are depicted. **d,** Same as in b, with a zoom in selected positive regions, and DBSCAN spatial clustering detection for extraction of cluster number (**e**) (P=0.023 by t-test) and molecule density (**f**) (P=4.6e-07 by t-test). Dashed line defines the cell boundary. **g,** Representative total internal reflection fluorescence (TIRF) microscopy of mCherry_talin1, (**h**) mCherry_paxillin and corresponding phalloidin-stained actin (**g,h**) in POU5F1B- and LacZ-overexpressing SW480 cells. **i, j,** Quantification of adhesion distance to the cell center (0=periphery, 1=center) (talin 1 P=0.01; paxillin P<0.0001 by t-test) and relative frequency signal from images in g-h. **k,** Mean actin radial intensity +/- standard deviation calculated from the cell periphery (0) to the center (1) from images in g-h. Six technical replicates from 3 biological replicates are represented in i-j-k. **l,** Representative images from experimental dissociation kinetics of PA-GFP_talin1 from mCherry_paxilin-positive FAs and **n,** photoactivable(PA)-GFP_paxilin from mCherry_talin1-positive FAs in the same cells as in g-h. **m, o,** Dissociation kinetics plots and box plot of the half-lives (talin 1 P=0.0008; paxillin P=0.028 by t-test) from images in l-n. Data from 4 independent experiments are represented in m-o. *p ≤ 0.05; **p < 0.01; ***p < 0.001; ****p < 0.0001.

Confocal microscopy images revealed a sharp peripheral ring-like arrangement of p-Y^397^FAK-positive foci in wild-type, POU5F1B-expressing LS1034 cell colonies, contrasting with a more diffuse pattern in their knockdown counterparts (Supp. Fig. 4C, 4D). We confirmed these results by using total internal reflection fluorescence (TIRF) microscopy, which allows the selective visualization of cellular structures very close to the glass coverslip. We analysed the distribution of talin 1 (Fig. 4G), an integrin adaptor protein shown to activate and to cluster integrins^29^, and paxillin (Fig. 4H), which together with FAK is recruited to nascent adhesions at the cell periphery^30^. Measuring the distance of these adhesion structures from the cell periphery to the cell center revealed that POU5F1B-expressing SW480 cells had a more peripheral distribution of talin 1 (Fig. 4I) and paxillin (Fig. 4J) compared with SW480 LacZ control cells, where adhesions were more evenly distributed throughout the cell (Fig. 4I, 4J). This peripheral localization of FA components in POU5F1B-expressing cells coincided with the accumulation of actin filaments close to the cell edges, compared to control cells where they were more diffusely spread (Fig. 4K).

Cell movement depends on the assembly and disassembly (turnover) of FAs, which can be used by cancer cells to enhance metastatic dissemination^31^. We previously demonstrated through mouse xenotransplantation experiments that POU5F1B stimulates the formation of metastases by human colorectal cancer cells^12^. We hypothesized that the increased number, size and density of FAs (Fig. 4B-F) and their peripheral redistribution (Fig. 4G-J) in POU5F1B-expressing cells might be accompanied by a faster turnover of their components, which could facilitate cell mobility. To analyse the stability of adhesion structures, we quantified the dissociation rate of Talin1 from FAs via photoactivation experiments^32^. Talin1 was C-terminally tagged with photoactivable GFP, and co-expressed with mCherry-tagged paxillin to identify FAs, in which Talin1 molecules were subsequently photoactivated to study their dissociation rate. The fluorescence loss after photoactivation (FLAP) was recorded for 3 minutes and adhesion stability was assessed by the fluorescence decay in the paxillin-labeled FAs. This analysis showed that POU5F1B expression reduced the stability of Talin1, diminishing its half-life in focal adhesions by 2.4-fold compared to that measured in LacZ control SW480 cells (Fig. 4L-M). Similar results were observed when mcherry-Talin1 labelled adhesion sites were studied for the dissociation rate of photoactivable-GFP labelled paxillin (Fig. 4N-O), a protein we found enriched in DRMs of POU5F1B-expressing cells (Fig. 3K).

Thus, our data indicate that POU5F1B-expressing cells harbour more numerous and larger Y^397^FAK positive adhesive units accumulating in the cell periphery and containing more and denser Y^397^FAK nanoclusters. In addition, these adhesion structures display a more dynamic exchange of integrin adapter proteins, suggesting an increased ability to turn over adhesion structures towards enhanced cell mobility.

### POU5F1B drives spontaneous cell invasion in 3D

Cell migration is associated with FA and integrin turnover. The accelerated turnover of FAs observed in POU5F1B expressing cells, led us to evaluate cancer cell invasion using an *in vitro* three-dimensional (3D) assay. SW480 cell spheroids were embedded in a collagen-I gel and imaged by confocal microscopy 5 days after seeding (Fig. 5A). POU5F1B-overexpressing SW480 cells presented higher invasion capacity than cells expressing the GFP control or the K135E ubiquitination- or C185S-C198S acylation-defective POU5F1B mutants (Fig. 5B, 5C). We next used a 2D culture system that enables to monitor dissociation of individual cancer cells from 3D CRC spheroids (Fig. 5D). Spheroids were plated onto collagen-I coated dishes, as described^34^. Strikingly, WT POU5F1B-expressing SW480 cells displayed earlier first-cell dissociation times than LacZ-expressing control cells (Fig. 5E, 5F). This accelerated dissociation was abolished by the K135E or K182T mutations, indicating that POU5F1B cytoplasmic localization is necessary for this phenotype. Dissociation of individual cancer cells from 3D CRC spheroids has previously been shown to depend on the viscosity of the medium^33^. We thus switched from normal (0.77 cP) to high viscosity (8cP, achieved with 0.6% of methylcellulose) medium, which mimics the viscosity of body fluids in certain pathological situations^35^. Elevated viscosity accelerated the dissociation of individual cancer cells from 3D CRC spheroids (Fig. 5E, 5F), as previously described^34^. Yet, POU5F1B-expressing SW480 cells still displayed earlier first-cell dissociation times than LacZ-expressing controls. Thus POU5F1B induces an increase in cell dissemination from 3D tumor spheroids irrespective of mechanical cues. We further observed that the spheroid initiation capacity of POU5F1B-overexpressing SW480 cells, measured after plating of single cells (Fig. 5D), was greater than that of their K135E-, K182T, or LacZ-overexpressing counterparts (Fig. 5G). Altogether these experiments confirm that POU5F1B-expressing cells are more dynamic, consistent with their higher metastatic potential.

**Figure 5.**
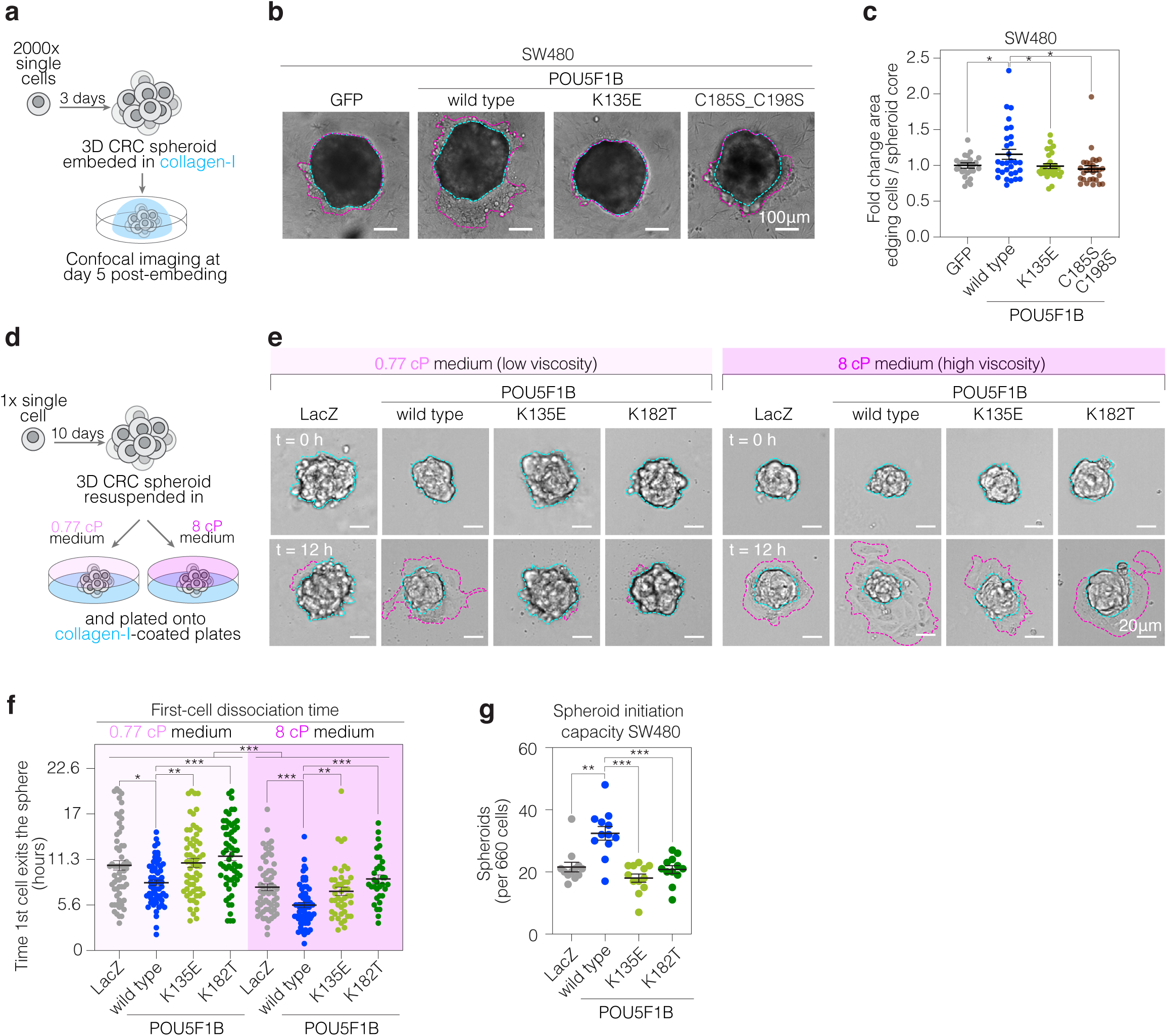
POU5F1B drives spontaneous cell invasion in 3D. **a,** Scheme ilustrating the preparation of 3D CRC spheroids embeded in collagen-I. **b,** Representative reflection contrast microscopy images of GFP-, POU5F1B-, POU5F1B-K135E-, POU5F1B-C185S_C198S-expressing SW480 spheroids. **c**, Fold change area of invading cells (in fucsia panel b) versus area spheroid core (in turquoise panel b) from spheroids in b (POU5F1B wild type vs. GFP P=0.031; vs. K135E P=0.037; vs. C185S_C198S P=0.017 by Wilcoxon test). **d,** Scheme ilustrating the preparation of 3D CRC spheroids plated onto collagen-I-coated plates and resuspended in 0.77 cP (low visocsity) and 8 cP (high viscosity) medium. **e,** Representative bright field confocal images from cells disseminating (fucsia contour) from LacZ-, POU5F1B-, POU5F1B-K135E-, POU5F1B-K182T-expressing SW480 3D spheroids (turquoise contour) cultured in low viscosity and high viscosity medium. **f,** The time required from the first cell to dissociate from each spheroid in panel e (n=60) from 3 experiments in low and high viscosity mediums (0.77 cP vs. 8 cP: LacZ vs. LacZ P=0.0009; POU5F1B vs. POU5F1B P= 8.57e-08; K135E vs. K135E P=9.57e-06; K182T vs. K182T P=0.0006. 0.77 cP experiment: POU5F1B vs. LacZ P=0.018; vs. K135E P=0.001; vs. K182T P=5.04e-06; 8 cP experiment: POU5F1B vs. LacZ P=0.0001; vs. K135E P=0.005; vs. K182T 1.57e-06 by Wilcoxin test). **g,** Number of spheroids grown from 660 LacZ-, POU5F1B-, POU5F1B-K135E-, POU5F1B-K182T-expressing SW480 single cells (POU5F1B wild type vs. LacZ P=0.003; vs. K135E P=0.0002; vs. K182T P=0.0006 by Wilcoxon test). * pvalue<0.05, ** pvalue<0.01, *** pvalue<0.001.

### Loss of ROCK activity abrogates POU5F1B function

POU5F1B is not endowed with enzymatic activity, and thus could be presumed non-druggable. However, approaches aimed at triggering the proteasomal degradation of pathogenic effectors have changed this perspective, as exemplified by the thalidomide-, lenalidomide-, or pomalidomide-induced degradation of IKZF1 and IKZF3^36–39^ for the treatment of multiple myeloma^40^. To explore this avenue, we searched for direct and indirect POU5F1B degraders using a recently described gain-of-signal cell-based assay^41^. For this, we established SW480 cell derivatives expressing POU5F1B fused to deoxycytidine kinase (DCK), an enzyme that converts the non-natural nucleoside 2-bromovinyldeoxyuridine (BVdU) into a poison^42^, together with green fluorescence protein (GFP) (Fig. 6A). Parental SW480 cells or derivatives expressing wild type POU5F1B or unfused DCK served as controls (Supp. Fig. 5A, 5B). We first confirmed that SW480 cells expressing DCK, either in its native form or fused to POU5F1B, were more sensitive to BVdU than unmodified SW480 cells or cells expressing wild-type POU5F1B (Supp. Fig. 5B). We then exposed POU5F1B-DCK cells in duplicate to a library of 5,758 known bioactive compounds, ∼25% of them FDA approved, for 24 hours before adding 200 µM BVdU to the culture in order to kill all cells but those in which degradation of the POU5F1B-DCK fusion protein had been induced by the pre-treatment (Fig. 6B). This first step of our screen singled out 34 compounds with a Z-score greater than the mean of the negative controls plus three standard deviations (Fig. 6C). These were then used to perform a counter-screen with dose response curves in both POU5F1B-DCK- and DCK-expressing SW480 cells. At the outset of this second step, only two drugs were found to protect POU5F1B-DCK-but not DCK-expressing cells from BVdU-induced death: capivasertib (or AZD5363) and OXA-06, which both are Rho kinase (ROCK) inhibitors (Fig. 6D). Importantly, the two compounds exerted negligible cytotoxicity on unmodified SW480 cells (Fig. 6D).

**Figure 6.**
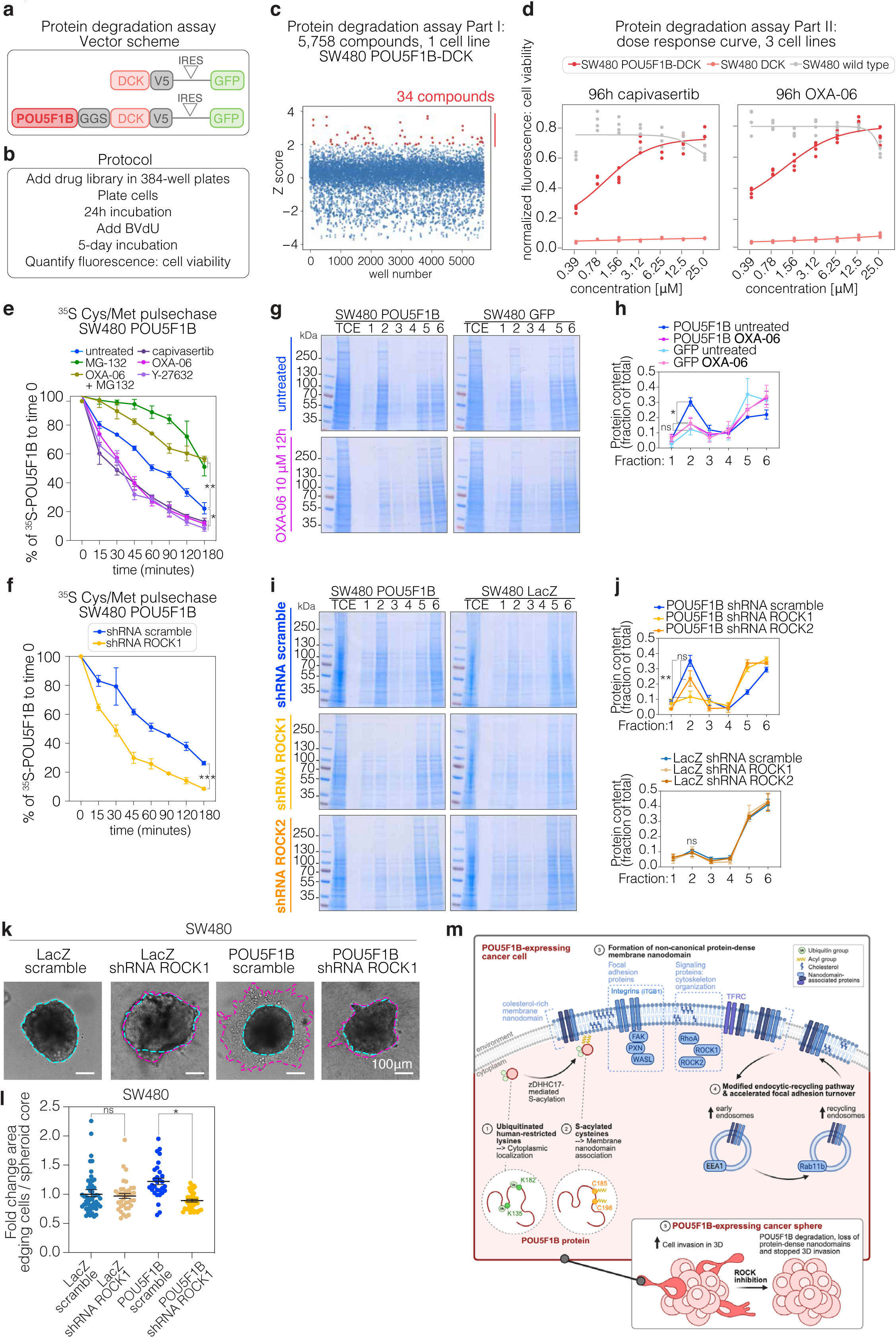
Loss of ROCK activity abrogates POU5F1B function. **a,** Vector scheme. DCK, deoxycytidine kinase; GGS, Gly-Gly-Ser spacer; IRES, internal ribosomal entry site. **b,** Protein degradation assay protocol. **c,** Z-distribution of GFP fluorescence of SW480 POU5F1B-DCK cells screened with the 5,758 known bioactive compounds library, and treated with 200uM BvdU 24h later. GFP signal was measured 48h post BvdU treatment. Colored in red, 34 compounds with a Z-score greater than the mean of the negative controls plus three standard deviations **d,** Dose response curves in wild type, POU5F1B-DCK-, and DCK-expressing SW480 cells treated with increasing concentrations of capivasertib and OXA-06 ROCK inhibitors. **e,** Pulse-chase experiment with 35SCys/Met in POU5F1B-expressing SW480 cells untreated or treated with MG-132 (10uM), OXA-06 (20uM), OXA-06 & MG-132, capivasertib (20uM), and Y-27632 (10uM). Drugs were added at time 0 and POU5F1B-HA protein was IP-ed from harvested cell pellets at indicated times (at time 180, untreated vs. MG-132 P=0.004; vs. OXA-06 P=0.026; vs. OXA-06 & MG-132 P=0.0011; vs. capivasertib P=0.03; vs. Y-27632 P=0.01 by t-test). **f,** Same as in (e) using SW480 POU5F1B cells expressing sh-scramble or sh-ROCK1 (P=0.0001 by t-test). **g,** Isolation of detergent-resistant membranes (DRMs) from POU5F1B-HA-wild-type-, and GFP-overexpressing SW480 cells untreated or treated with OXA-06 10 μM, fractions were run on SDS-PAGE and gels were stained with Coomassie. 1-3 correspond to insoluble fractions, 4-6 to soluble fractions, DRMs being traditionally found in fraction 2. **h,** Quantification of coomassie staining in (g) (POU5F1B untreated vs. POU5F1B OXA-06 P=0.02; GFP untreated vs. GFP OXA-06 P=0.95 by t-test). **i,** Same as in (g) from SW480 POU5F1B or SW480 GFP cells expressing either sh-scramble or sh-ROCK1/2. **j,** Quantification of coomassie staining in (i) (POU5F1B shRNA scramble vs. shROCK1 P=0.007; vs. shROCK2 P=0.13; LacZ shRNA scramble vs. shROCK1 P=0.77; vs shROCK2 P=0.64 by t-test). **k,** Representative reflection contrast microscopy images of 3D SW480 spheroids embeded in collagen-I, as in Figure 5a/b. **l,** Fold change area of invading cells (in fucsia panel k) versus area spheroid core (in blue panel k) from LacZ scramble-, LacZ shRNA ROCK1-, POU5F1B scramble-, and POU5F1B shRNA ROCK1-expressing SW480 spheroids (LacZ scramble vs. shROCK1 P=0.99; POU5F1B scramble vs. shROCK1 P=1.32e-06 by Wilcoxon test). ns not significant, * pvalue<0.05, ** pvalue<0.01. **m,** Model. (1) Ubiquitination of K135 and K182 human-restricted lysines retains POU5F1B protein in the cytoplasm of cancer cells. (2) S-acylation of C185 and C198 cysteines by ZDHHC17 S-acyltransferase associates POU5F1B to cholesterol-rich membrane nanodomains. (3) POU5F1B induces the formation of transferrin receptor(TFRC)-containing non-canonical protein-dense membrane nanodomains, enriched in focal adhesion (integrins -ITGB1-, FAK, PXN, WASL) and signaling proteins involved in cytoskeleton organization (RhoA, ROCK1, ROCK2). (4) POU5F1B-expressing cells present modified endocytic-recycling pathway and accelerated turnover of focal adhesion structures. Higher number of EEA1-positive early endosomes and Rab11b-positive recycling endosomes is representative of these cells. (5) POU5F1B-expressing cancer spheres present augmented cell invasiveness in 3D. ROCK inhibition triggers POU5F1B degradation, loss of protein-dense nanodomains and stops 3D invasion. Created with Biorender.com.

We validated these results by performing pulse-chase experiments with ^35^SCys/Met in SW480 cells. Our results revealed that the half-life of POU5F1B was 60 min in untreated SW480 cells and increased to 160 min in the presence of the proteasome inhibitor MG132 (Fig. 6E), indicating that it is degraded by the proteasome. In sharp contrast, the half-life of POU5F1B dropped to 30 min when cells were treated with capivasertib, OXA-06 or Y-27632, another ROCK inhibitor, yet was restored to 160 min in the combined presence of OXA-06 and MG132 (Fig. 6E). This result indicates that ROCK inhibition triggers POU5F1B proteasomal degradation, and in reverse that ROCK stabilizes POU5F1B. The half-life of POU5F1B was similarly reduced upon dox-inducible ROCK1 knockdown (Fig. 6F, Supp. Fig. 5C). Furthermore, upon 24 hours of either OXA-06 treatment (Fig. 6G, 6H) or ROCK1 downregulation (Fig. 6I, 6J), the overall protein content of DRMs (fraction 2) decreased in POU5F1B-overexpressing but not control SW480 cells (Fig. 6G, 6H, 6I, 6J). Interestingly, a stronger effect was observed upon ROCK1 than ROCK2 downregulation, indicating that POU5F1B is more dependent on ROCK1 in these cells (Fig. 6I, 6J). Consistently, downregulation of ROCK1 lowered the amount of POU5F1B associated with DRMs (Supp. Fig. 5D). Noteworthy, the remaining amounts of ROCK1 were also no longer found in DRMS in this setting (Supp. Fig. 5D). Finally, ROCK1 downregulation decreased the invasion capacity of POU5F1B-overexpressing SW480 cells in the 3D spheroid invasion assay, whereas it had no effect on control cells (Fig. 6K, 6L).

## Discussion

Our work demonstrates that the product of the human ortholog of the hominoid-restricted *POU5F1B* gene is endowed with unique properties dependent on sequential ubiquitination and S-acylation. The first of these post-translational modifications targets residues specific to the human protein, meaning that *POU5F1B* represents a human-unique cancer-promoting gene, to our knowledge the first described to date. While another human-specific retrogene, *NANOGP8*, has also been found overexpressed in some cancers, its product is similar in sequence and function to that of its parental *NANOG* gene, which encodes a well characterized stemness-promoting transcription factor^43^. As well, although the onco-exaptation of human-restricted TE integrants could theoretically result in the making of oncoproteins with unique properties, in examples investigated so far the function of the product made from these TE-driven aberrant transcripts was found to be similar to that of their native counterparts, for example the AluJb-LIN28B fusion protein that carries the same function as LIN28B despite a 22 residue-long N-terminal extension^44^.

Our biochemical and functional studies demonstrate that POU5F1B exerts its pro-oncogenic action through an unprecedented mechanism: the formation of non-canonical protein-enriched membrane nanodomains. Membrane nanodomains are known to play important role in signal transduction through their capacity to act as protein concentrating platforms^45^. Malignant cells exploit these platforms to initiate oncogenic signaling and promote tumor progression^46^. Although many studies have focused on the consequences of nanodomain formation in cancer biology^46^, little is known about the intracellular mechanisms fostering their assembly in this context. Membrane nanodomains are known to be enhanced by extracellular cues such as cell-to-cell contacts, as in the setting of the immune synapse^47^, or by ECM-to-cell contacts, as when migrating cells concentrate adhesion proteins in response to increased ECM stiffness^48^. The ECM component agrin had been found to foster the clustering of membrane nanodomains in both immune and nervous synapses^49^. Here we identify POU5F1B as stimulating the remodeling of membrane nanodomains from inside the cells, profoundly remodeling their protein content. We discovered several signaling molecules (e.g. SMAD2, SMAD4, MAPK1, FER) to be enriched in DRMs of POU5F1B positive cells, and we previously reported that POU5F1B co-immunoprecipitates with a number of signaling proteins (e.g. ERBB2, MST1R, PRKCA)^12^. DRMs of POU5F1B-expressing cells were also enriched with FA-building proteins, which organized in high-density FA structures that displayed increased rates of turnover and colocalized with a peripheral organization of actin filaments. We also observed the accumulation of RhoA, ROCK1 and ROCK2 in these altered membrane nanodomains and discovered that POU5F1B-triggered phenotypes depend on the activity of ROCK, the blockade or depletion of which destabilized the oncoprotein. It has been previously shown that activated RhoA increases myosin-dependent contraction of the actin cytoskeleton^50^, leading to intracellular tension and increased turnover of integrins in FAs^51^, a prerequisite for cell migration. This could partially explain why POU5F1B-expressing cells are capable of migrating faster out of a spheroid even in the absence of an external physical stimulation (e.g. increase of medium viscosity). Although we observed that POU5F1B positive cells transform membrane nanodomains into denser signaling and adhesion platforms, we do not know yet whether signaling and adhesion molecules are found within the same or distinct sets of nanodomains.

The continuous endocytosis and recycling to the plasma membrane of focal adhesions building blocks is an essential component of cell movement^52^, and cholesterol-sphingolipid rich membrane nanodomains are implicated in endocytosis^53^, sorting and recycling^54–56^ processes. Accordingly, we describe here that POU5F1B-expressing cells have membrane nanodomains enriched in endocytic proteins, and display an enhanced vesicular endocytic-recycling pathway as well as a higher turnover of focal-adhesion related molecules. We previously observed these cells present increased levels of caveolin-1 and other components of caveolae^12^, which are the invaginated form of plasma membrane nanodomains^57^ that associate with the activation of the contractile actin cytoskeleton and favor cell migration^58^. It suggests that POU5F1B-induced alterations in endocytosis and recycling might proceed through caveolae.

The actin-myosin cytoskeleton and its regulatory axis Rho GTPase-ROCK proteins are used by cancer cells to migrate and metastasize^59,60^. ROCK proteins regulate cell contractility, actin cytoskeleton organization and FA formation, which we found to be enhanced by POU5F1B. Future studies will address whether POU5F1B is a direct target of ROCK or whether it is stabilized by interacting with a ROCK-phosphorylated partner. Our pharmacological and genetic data concur to demonstrate that ROCK activity is essential to the stability hence the function of POU5F1B. Indeed, blocking or depleting the kinase triggers POU5F1B proteasomal degradation, with *ROCK1* knockdown abrogating the increased cell invasiveness induced by the retrogene product in a 3D spheroid assay. While ROCK inhibitors have so far been seldom used to treat cancer^61–63^, our data warrant exploring their therapeutic potential under the lens of POU5F1B expression, whether as a biomarker for patient stratification or to guide the selection of specific inhibitors. *POU5F1B* emerged as the result of a retrotransposition event that occurred some 12 to 15 million years ago. The physiological roles of its human product and the nature of the pressures that drove its selection during the last 5 to 7 million years, that is, after the human-chimpanzee split, remain to be identified. Irrespectively, as the product of a retrogene expressed in cancer by aberrant activation of upstream TE-embedded regulatory sequences, POU5F1B provides a multifaceted illustration of the prominent role played by transposable elements in conferring an often-underestimated degree of species-specificity to the biology of higher vertebrates.

## Acknowledgements

We thank the members of Alexandre Reymond’s and Didier Trono’s laboratories for stimulating discussions. We gratefully acknowledge the Proteomics Core Facility (PCF), and the Bioimaging and Optics Platform (BIOP) at EPFL, as well as the Imaging Platform ACCESS and the imaging core facility (CMU) at the University of Geneva for their skilful assistance. We are particularly grateful to David Root (Genetic Perturbation Platform, Broad Institute of Harvard and MIT) for helping us set up the screen for inducers of POU5F1B degradation. This work was supported by grants from the European Research Council (Transpos-X, No. 694658), the Personalized Health and Related Technologies (PHRT-508) program, the Swiss National Science Foundation (310030-152879, 310030B-173337, 310030-18803, 320030-227571, 310030-192613 and 1310030-219201), and the Novartis foundation No. 532316 to D.T., the Swiss Cancer Research Foundation (KFS-4968-02-2020 and KFS-5823-02-2023) to D.T. and L.S.R., the UNIL Tremplin grant to L.S.R., and grants from the Swiss National Science Foundation to B.W.H. (320030-228395) and GVDG. G.E.F. acknowledges the support of the H2020 - UE Framework Programme for Research & Innovation (2014-2020); ERC-2017-CoG; InCell; Project number 773091, and the EPFL Centre for Imaging, grant number 21645. The authors acknowledge the use of <u>BioRender.com</u> for the creation of the graphical abstract and the model depicted in Figure 6M.

## Methods

### Cell culture

LS1034 and SW480 cell lines were obtained from the American Type Culture Collection (ATCC) and maintained in RPMI 1640 (Gibco) and L15 medium (Sigma) respectively, being supplemented with 10% FCS (Bioconcept 2-01F36-I). For lentiviral vector production, 293T cells were cultured in DMEM supplemented with 10% FBS with 100 IU ml^-1^ penicillin, 100 ug ml^-1^ streptomycin, and 26 µg ml^-1^ glutamine (Corning 30-009-CI) at 37°C in a humidified atmosphere of 5% CO_2_. All cells tested negative for mycoplasma.

### Overexpression and shRNA vectors

For the POU5F1B-expressing vector, genomic DNA from DLD1 cells (ATCC® CCL-221™) was PCR amplified with the POU5F1B primers 5’-CACCATGGCGGGACACCTGGCTTCGGATTTC-3’, 5’-GTTTGAATGCATGGGAGAGCCCAG-3’, cloned into a pENTR TOPO donor vector (Thermo Fisher), that was recombined with the doxycycline-inducible lentiviral destination vector pSin-TRE-3xHA-puro. For the DCK-based positive selection assay, we kindly received from William Kaelin’s laboratory the pLX304-gate-DCK-V5-IRES-GFP plasmid, in which we Gateway cloned *POU5F1B*. Short hairpin RNA (shRNA) were designed with the help of the i-Score Designer website, and adapted to the miR-E shRNA structure and cloned into the LT3GEPIR pRRL backbone following Fellmann C et. al. instructions^64^: shRNA3 5’-TGCTGTTGACAGTGAGCG**CACAGGTGATTATGATTTAAAG**TAGTGAAGCCACAGATGT A**CTTTAAATCATAATCACCTGTG**TGCCTACTGCCTCGGA-3’; shRNA5 5’-TGCTGTTGACAGTGAGCG**CACATTCAGTCAACATTTAATG**TAGTGAAGCCACAGATGT A**CATTAAATGTTGACTGAATGTG**TGCCTACTGCCTCGGA-3’) with a scrambled shRNA serving as negative control 5’-TGCTGTTGACAGTGAGCG**CAACAAGATGAAGAGCACCAAG**TAGTGAAGCCACAGATG TA**CTTGGTGCTCTTCATCTTGTTG**TGCCTACTGCCTCGGA-3’.

### Lentivirus production and stable gene expression

Lentiviral particles produced as described at http://tronolab.epfl.ch were used to transduce SW480 and LS1034 cells in 6-well plates at multiplicity of infection 1, before selection in 1 μg ml^-1^ puromycin for 7 days. POU5F1B, GFP and lacZ overexpression in SW480 cells was verified, after three days of 500 ng ml^-1^ doxycycline activation, by western blot with HRP-conjugated anti-HA antibody (clone 3F10 Roche 12013819001, 1:1,000); and *POU5F1B* downregulation upon shRNA transduction in LS1034 was analyzed by quantitative real-time PCR using P3 pair of primers^12^. Constitutive expression of POU5F1B-DCK and DCK proteins was verified by western blot with anti-V5 antibody.

### Immunofluorescence

Cell lines were plated on glass coverslips into 24-well plates and cultured for 3 days up to 70% confluence in 500 ng ml^-1^ doxycycline-containing medium. Growth medium was refreshed (1 ml) prior a 15 min fixation with 1 ml 8% paraformaldehyde (PFA) added dropwise, to achieve a final concentration of 4% PFA. Cells were washed three times with PBS, permeabilized with 0.05% saponin PBS for 20 min, and blocked with 1% BSA 0.05% saponin PBS for 30 min before incubation with anti-HA (bioLegend, Covance catalog #MMS-101P, 1:1,000) and anti-lamin-B1 (abcam ab16048, 1:1,000) antibodies in 1% BSA 0.05% saponin PBS 2 h at room temperature (RT) under agitation. Phospho-FAK Tyr397 (Thermofisher 700255, 1:200) antibody was incubated overnight at 4 degrees in blocking solution without saponin. Samples were washed three times with PBS and incubated with Alexa 647-conjugated (A647) anti-mouse antibody (1:1,000) and A568 anti-rabbit antibody (1:1,000) in 1% BSA PBS for 40 min at RT. Three final washes were performed before mounting the slides in Vectashield with DAPI (Vector Laboratories). Images were acquired on a ZEISS LSM 700 confocal microscopy and analyzed with Fiji software.

### Immunoprecipiation and western blot

For immunoprecipitation and western blot, cells were lysed on ice for 30 min with lysis buffer (500 mM Tris–HCl pH 7.4, 2 mM benzamidine, 10 mM NaF, 20 mM EDTA, 0.5% NP40 and a protease inhibitor cocktail (Roche, CH)). Samples were sonicated using a probe sonicator (two times 5 seconds at 30% amplitude). The lysate was then clarified by centrifugation at 4 °C for 5 min at 5000 rpm and incubated overnight on a rotating wheel at 4 °C with 50 µl of HA coupled agarose beads (Sigma 11867423001). The beads were then washed 3× with lysis buffer before adding 4× Sample Buffer including beta-mercaptoethanol. Similar elution volumes and protein quantities for the IPs and the inputs, respectively, were submitted to SDS-PAGE and analyzed by immunoblotting using HRP-conjugated anti-HA (clone 3F10 Roche 12013819001, 1:1,000) and anti-Ubiquitin antibody. ***In vitro* proliferation assays.** Cell proliferation was determined with PrestoBlue™ reagent (A13262, Invitrogen) for 6 days.

### Isolation of detergent-resistant membranes (DRMs)

Approximately 1 × 10^7^ cells were resuspended in 0.5 ml cold TNE buffer (25 mMTris-HCl, pH 7.5, 150 mM NaCl, 5 mM EDTA, and 1% Triton X-100; Surfact-Amps, ThermoFisher) with a tablet of protease inhibitors (Roche). Membranes were solubilized in a rotating wheel at 4 °C for 30 min. DRMs were isolated using an Optiprep^TM^ gradient ^65^: the cell lysate was adjusted to 40% Optiprep^TM^, loaded at the bottom of a TLS.55 Beckman tube, overlaid with 600 μl of 30% Optiprep^TM^ and 600 μl of TNE, and centrifuged for 2 h at 55,000 rpm at 4 °C. Six fractions of 400 μl were collected from top to bottom. DRMs were found in fraction 2. Equal volumes from each fraction were analyzed by SDS-PAGE and western blot analysis using HRP-conjugated anti-HA, caveolin1 (Santa Cruz sc-894, 1:500) and transferrin receptor (ThermoFisher 13-6800, 1:1,000) antibodies.

### Acyl-RAC capture assay

Protein S-palmitoylation was assessed by the Acyl-RAC assay as previously described^66^ with some modifications. Cells or supernatants were lysed in 400 ml buffer (0.5% Triton-X100, 25 mM HEPES, 25 mM NaCl, 1 mM EDTA, pH 7.4,and protease inhibitor cocktail). Cell lysis were incubated 30 min at RT with 10mM TCEP. Then, 200 ml of blocking buffer (100 mMHEPES, 1 mM EDTA, 87.5 mM SDS, and 1.5% [v/v] methyl methanethiosulfonate (MMTS)) was added to the lysates and incubated for 4 h at 40°C to block free the SH groups with MMTS. Proteins were acetone precipitated and resuspended in buffer (100 mMHEPES, 1 mM EDTA, 35 mM SDS). For treatment with hydroxylamine (NH2OH) and capture by Thiopropyl Sepharose beads, 2 M of hydroxylamine was added together with the beads (previously activated for 15 min with water) to a final concentration of 0.5 M of hydroxylamine and 10% (w/v) beads. As a negative control, 2 M Tris was used instead of hydroxylamine. These samples were then incubated overnight at room temperature on a rotating wheel. After washes, the proteins were eluted from the beads by incubation in 40 µl SDS sample buffer with ß-mercaptoethanol for 5 min at 95°C. Finally, samples were separated by SDS-PAGE and analyzed by Western blot. A fraction of the cell lysate (total cell extract) was saved as the input.

### Radiolabeling 3H-palmitic acid incorporation

To follow S-acylation, lentiviral vector transduced and 96h doxycycline activated cells were incubated 1 hour in L15 medium without serum, followed by 2 hours at 37°C in L15 without serum with 200 uCi /ml 3 H palmitic acid, washed with cold PBS, and lysed with RIPA buffer prior overnight anti-HA immunoprecipitation. Beads were incubated 5 min at 90°C in reducing sample buffer prior to SDS-PAGE. Immunoprecipitates were split into two, run on 4-20% gels and analyzed either by autoradiography (3 H-palmitate) after fixation (25% isopropanol, 65% H2O, 10% acetic acid), incubation 30 min in enhancer Amplify NAMP100, and drying; or western blot.

### Transfection of siRNA

SW480 cells were transfected with siRNA (Qiagen, Supplementary Table S1) (15 pmol/ 9.6cm2 plate) using Lipofectamine RNAiMAX transfection reagent. Cells were harvested 72h post transfection and whole cell protein lysates were analyzed by western blot.

### Scanning ion conductance microscopy (SICM) imaging, data processing and analysis

Images from live SW480 cells were taken in L-15 (Leibovitz) medium, at 37°C and saturated humidity. Imaging was performed with bias voltages of 200 to 300mv, 2 to 4nA ion currents and typical pipette tip inner diameters of 100nm. A detailed description of the homebuilt microscope platform has been published here^24^. SICM images were processed using Gwyddion^67^. Image defects with the maximum width of 1 pixel were removed by filling-in the neighboring line. The cells were marked by implementing height thresholding, with the height data of the images leveled by shifting the mean height value of the substrate part to zero. To analyze the roughness and height of the cell, we separated the small structure on the cell membrane surface from the cell topography by using frequency split or two polynomial degree leveling. The cell bodies were masked by shrinking the mask of the cells. The root mean square roughness of a cell body and boarder were calculated from the high frequency image, and cell height was extracted by using the max value of the low frequency image. All SICM topography data were normalized by 13 µm and exported in 16-bit Portable Network Graphics format with gray and customized color bar. We used the open-source 3D rendering tool Blender 3D to generate the three-dimensional images of the cells. The corresponding SICM height data and colored data were imported and used as a height map and projected color on the topography.

### Cell surface protein studies

Surface biotinylation assays were performed as previously described^68^. Briefly, cells were shifted to ice and incubated for 30 min with cold biotinylation solution (0.2 mg/ml sulfo-NHS-SS-biotin (Pierce) diluted in PBS -0.17-mg/mL final concentration-), then quenched three times with cold 100 mM NH4Cl, and lysed in IP buffer. The lysate was immunoprecipitated overnight at 4°C with prewashed streptavidin-coated beads (Sigma S1638), beads were washed with 1 mL 1% SDS, 150 mM NaCl, 1 mM EDTA in 50 mM HEPES pH 7.4 (×3) and 50 mM HEPES pH 8.0 (×2), and proteins were eluted from beads with 50 µL of 50 mM HEPES pH 8.0 containing 50-100 mM TCEP for 1 h. Peptides containing supernatants were digested overnight at 37 ⁰C in the thermoshaker with 1-2 µL Trypsin (20 µg /100 µL) in 50 mM HEPES pH 8. For LC MS/MS analysis, peptides were resuspended and separated by reversed-phase chromatography on a Dionex Ultimate 3000 RSLC nanoUPLC system in-line connected with an Orbitrap Lumos Fusion Mass-Spectrometer. Database search was performed using MaxQuant 1.6.10.43^69^ against concatenated database consisting of the UniProt human database (Uniprot release 2019_06, 74468 sequences) and common fetal bovine serum protein^70^. Raw spectral counts extracted with MaxQuant were subjected to log2 transformation, and any missing values were imputed using the ‘min’ function available in the DEP package^71^. Proteins that exhibited fewer than 10 log2-transformed total spectral counts across all samples were excluded for further analysis. To identify proteins undergoing significant changes between conditions of interest, a statistical linear model with interactions was applied using the limma R package^72^. Surface-associated proteome candidates were selected when the three experimental replicates showed p value<0.05 (outlier detection test as computed by MaxQuant), high intensity (above quantile 25%), and fold changes in the same direction for the two shRNAs.

### LC-MS/MS analysis of DRM fractions

DRM fraction 2 (F2) and equal volume mixes of fractions 4, 5, and 6 (F456) from shRNA-scramble, shRNA3 and shRNA5-transduced LS1034 cells were analyzed by Liquid Chromatography with tandem mass spectrometry (LC-MS-MS). Each sample was digested by Filter Aided Sample Preparation (FASP)^73^ with minor modifications. Dithiothreitol (DTT) was replaced by Tris (2-carboxyethyl)phosphine (TCEP) as reducing agent and Iodoacetamide by Chloracetamide as alkylating agent. A combined proteolytic digestion was performed using Endoproteinase Lys-C and Trypsin. Peptides were desalted on SDB-RPS StageTips^74^ and dried down by vacuum centrifugation. Samples were then fractionated into 12 fractions using an Agilent OFFGEL 3100 system. Resulting fractions were desalted on SDB-RPS StageTips and dried by vacuum centrifugation. Samples were analyzed by LC-MS/MS as described above. Proteins that demonstrated notably different behavior between the ‘scramble’ and ‘sh’ conditions in F2 compared to their F456 counterparts, and exhibited a foldChange of less than 0.5 log2 in all ‘inputs’, were selected for further analysis.

### DAVID enrichment analysis

For DRM and surface associated-proteome data sets, we obtained a list of differentially expressed candidates that were selected as detailed above. Resulting candidate lists were subjected to GO and KEGG functional annotation chart using the online bioinformatics resource DAVID^25,26^.

### High resolution microscopy imaging

#### Buffers for super-resolution experiments

The following buffers were used for sample preparation and imaging:

- Buffer C+: 1× PBS, 500 mM NaCl and 0.05% Tween-20 (Sigma Aldrich cat: P9416-50ML)
- Buffer C: 1× PBS, 500 mM NaCl
- Antibody Incubation buffer: 1×PBS, 1 mM EDTA, 0.02% Tween-20, 0.05% NaN3, 2% BSA (Sigma Aldrich cat: A4503-10G), and 0.05 mg/ml sheared salmon sperm DNA (Thermo Fisher cat: 15632011)
- Blocking Buffer: 1×PBS, 1 mM EDTA, 0.02% Tween-20, 0.05% NaN3, 2% BSA (Sigma Aldrich cat: A4503-10G), 0.05% saponin and 0.05 mg/ml sheared salmon sperm DNA (Thermo Fisher cat: 15632011)
- Imaging solution: Buffer C+ supplemented with oxygen scavenger system (1x Trolox, PCA, PCD)

#### Oxygen scavenger system preparation

100× Trolox (Sigma Aldrich cat: 238813-5 G): 100 mg Trolox was added to 430 μl 100 % Methanol, 345 μl 1M NaOH, and 3.2 ml H2O. 40× PCA (Sigma Aldrich cat: 37580-25G-F): 154 mg PCA, 10 ml water and NaOH were mixed and pH was adjusted 9.0. 100× PCD (Sigma Aldrich cat: P8279): 9.3 mg PCD, 13.3 ml of buffer (100 mM Tris-HCl pH 8, 50 mM KCl, 1 mM EDTA, 50 % Glycerol).

#### Nanobody-DNA conjugation via single cysteine

Rabbit secondary nanobodies were purchased from Nanotag (cat: N2405) with a single ectopic cysteine at the C-terminus for site-specific and quantitative conjugation. The conjugation to DNA-PAINT docking site 7xR4 – ACACACACACACACACACA with a 5’ C-3 azide (Metabion) - was performed as previously described (cite: doi.org/10.1039/D0NR00227E). In brief, the buffer was exchanged to 1× PBS + 5 mM EDTA, pH 7.0 using Amicon centrifugal filters (10k MWCO) and free cysteines were reacted with a 20-fold molar excess of bifunctional maleimide-DBCO linker (Sigma Aldrich, cat: 760668) for 2-3 hours on ice. The unreacted linker was removed by buffer exchange to PBS using Amicon centrifugal filters. Azide-functionalized DNA was added with 3-5 molar excess to the DBCO-nanobody and reacted overnight at 4°C. Unconjugated nanobody and free azide-DNA were removed by anion exchange using an ÄKTA Pure liquid chromatography system equipped with a Resource Q 1 ml column. Nanobody-DNA concentration was adjusted to 5 µM (in 1xPBS, 50% glycerol, 0.05% NaN3) and stored at -20°C.

#### Super-resolution microscopy setup

Super-resolution DNA-PAINT imaging was performed on an inverted Eclipse Ti2 microscope (Nikon Instruments) with a Perfect Focus System unit, applying an objective-type TIRF configuration equipped with an oil-immersion objective (Nikon Instruments, Apo SR TIRF×100, NA 1.49, Oil). A 560-nm laser (MPB Communications, 1 W) was used for fluorophore excitation and coupled into the microscope via a Nikon manual TIRF module. The laser beam was passed through a cleanup filter (Chroma Technology, ZET561/10) and further coupled into the microscope objective using a beam splitter (Chroma Technology, ZT561rdc). Fluorescence was spectrally filtered with an emission filter (Chroma Technology, ET600/50m, and ET575lp) and detected on an sCMOS camera (Hamamatsu Fusion BT) without further magnification, resulting in an effective pixel size of 130 nm after 2×2 binning. TIR illumination was used for all measurements. The central 1152×1152 pixels (576×576 after binning) of the camera were used as the imaging field of view (FOV). Raw microscopy data was acquired using μManager (Version 2.0.1) (cite: doi: 10.14440/jbm.2014.36).

#### DNA-PAINT sample preparation and imaging

SW480 POU5F1B-overexpressing and GFP control cell lines were seeded with approximately 30K cells per cm2 in 8-well ibidi µ-slides (ibidi cat: 8080) with doxycycline supplemented media (500 ng/ml) for 96 h (replenishment with fresh media after 48h). Afterwards cells were fixed with 4% PFA, washed 4 times with PBS and permeabilized with 0.05% saponin in PBS for 15 min. Cells were then washed with PBS three times and blocked with Blocking buffer for 30 min. After 3 times washing with PBS 90 nm gold nanoparticles (1:2, Cytodiagnotics cat: G-90-100) were incubated onto the sample for 5 min followed by 3 times washing with PBS. Phospho-FAK (Tyr397) Recombinant Rabbit Monoclonal Antibody (31H5L17) (Thermo Fisher cat:700255) was diluted 1:200 and put onto the sample in antibody incubation buffer overnight. The next morning the sample was washed 4 times with PBS and a 1:200 dilution of rabbit secondary nanobody (7xR4) (Nanotag cat: N2405) was introduced to the sample in antibody incubation buffer (concentration of ∼50 nM) for ∼1 h. Finally, the sample was washed 4 times with PBS, once with buffer C and then incubated with imaging solution. For low resolution, high statistics experiments, the samples were supplemented with 500 pM R4 imager - TGTGTGT-Cy3B - (Metabion) containing imaging solution. The samples were mounted onto the microscope stage and 7.500 75 ms frames were acquired at a laser power of 30 mW (∼150 W/cm2). To ensure unbiased detection of highly expressing p-Y397FAK focal adhesion sites, FOV selection was done in brightfield. Three distinct samples per condition were imaged. For high resolution experiments the samples were supplemented with 100 pM R4 imager containing solution and 40.000 100 ms frames were acquired at 30 mW laser power. In order to select only highly expressing p-Y397FAK focal adhesion sites, FOV were prescreened with a high (500 pM) imager concentration and afterwards incubated with 100 pM imager for highest resolution imaging.

#### Single-molecule localization analysis

Raw fluorescence data was reconstructed using the Picasso software package (cite:doi.org/10.1038/nprot.2017.024) (the latest version is available at https://github.com/jungmannlab/picasso). Drift correction was performed using the AIM algorithm (cite: /doi.org/10.1126/sciadv.adm7765) with 90 nm gold nanoparticles (Cytodiagnotics cat: G-90-100) as fiducials for all experiments.

#### p-Y397FAK postprocessing analysis

For the low resolution, high statistics experiments, the 75 x 75 µm2 FOVs of POU5F1B-overexpressing and GFP control datasets were segmented for high intensity p-Y397FAK regions. First, gold nanoparticles were removed from localized and drift-corrected DNA-PAINT. The datasets were blurred using a 400 nm 2D bin size and 2.5 as gaussian blur in bin size units. The threshold for the masked region was set to 350 localizations for the p-Y397FAK regions and to 100 localizations for segmentation of the cell area (in a blurred bin), ensuring comparable segmentation of POU5F1B-overexpressing and GFP control samples. Area and density were calculated using the number of localizations inside the 2D binned masked regions.

For the high-resolution datasets, a 350 nm bin size 2D mask with 1.5 gaussian blur was generated for all datasets to mask the p-Y397FAK regions. The threshold was normalized to 0.15 times the brightest bin, which were generated using gold nanoparticles. After the masks were generated, gold was removed from the datasets. All localizations inside the masked regions were extracted and DBSCAN using 20 nm as search radius (approximately the label size of antibody and secondary nanobody) with minimal sample size of 40 localizations. With an imager concentration of 100 pM and 40.000 frames the expected number of binding events for R4 (on rate, kon of 27 * 106) (cite: doi.org/10.1038/s41592-020-0869-x) can be calculated. Using kon*cR4 = ξ, with c as concentration and ξ as influx rate, the influx rate of the measurements yields 0.0027 s-1 (cite: doi.org/10.1038/nmeth.3804). Since the total imaging time equals 40.000 times 100 ms, the total number of binding events in that duration is ∼10. Assuming a bright time of 0.4 s, a single binding event will result in 4 localizations for a single binding site, resulting in ∼40 localizations over the entire measurement. The area, localization numbers and densities for the DBSCAN clusters of POU5F1B-overexpressing and GFP control samples were then extracted on an individual cluster level for all datasets.

### Focal adhesion centrality analysis using total internal reflection fluorescence (TIRF) microscopy

LacZ-, and POU5F1B-expressing SW480 cells were transiently transfected with pcDNA3-mCherry-paxillin or pcDNA3-mCherry-talin1 using the JetPrime reagent (Polyplus) according to the manufacturer’s instructions. After transfection, cells were plated in home-made microscopy-compatible dishes and cultured for an additional 72 hours. Fixation was carried out using 2% paraformaldehyde (PFA) in PBS for 15 minutes at room temperature. Imaging was performed using Total Internal Reflection Fluorescence (TIRF) microscopy on a Nikon Eclipse Ti system equipped with Perfect Focus System (PFS) and a 100×/1.49 NA oil immersion objective, to visualize focal adhesion markers. Fluorescence images were processed in FIJI (ImageJ). Individual cell outlines were manually traced to define cell boundaries. A Euclidean distance map (EDM) was generated from each cell mask and the distance values were normalized by the maximum radial distance of the cell to produce a relative scale ranging from 0 (periphery) to 1 (center). Focal adhesions were detected with an intensity threshold, followed by region of interest (ROI) creation using the “Analyze Particles” function. These ROIs were overlaid onto the normalized distance maps, and the mean value within each ROI was computed, representing the average position of each adhesion relative to the cell center. Six cells were analyzed per condition in each experiment, with three independent experimental repeats.

### Actin centrality analysis using total internal reflection fluorescence (TIRF) microscopy

LacZ-, and POU5F1B-expressing SW480 cells were stained with Phalloidin-iFluor 647 (Cayman Chemical) to visualize actin and imaged using TIRF microscopy with a Nikon Eclipse Ti equipped with PFS and a 100x 1.49 NA objective. To quantify the spatial distribution of actin, we used a custom MATLAB pipeline to measure fluorescence intensity along radial lines from the cell centre to the periphery. In the first script, a manually defined ROI (drawn in Fiji and saved as a .roi file) was used to outline cell contours. Radial intensity profiles were then extracted from the centroid outward at 1° angular steps (360 directions per cell), with intensity values interpolated along each line. The distances were normalized from 0(periphery) to 1 (cell center), and the resulting raw and binned data (mean, SEM, and standard deviation) were exported to Excel for each cell. Subsequently, interpolated radial profiles from multiple cells and conditions were combined. Technical replicates (n = 6) were averaged for each biological replicate (N = 3), and the mean intensity profiles, along with SEM, were calculated across biological replicates for each condition. Curves were plotted as mean ± SEM across the normalized radial distance.

### Photoactivation experiments

Photoactivation-based experiments (FLAP). Image acquisition and analysis were performed as previously described^32^. Briefly, transfected SW480 cells were cultured overnight on coverslips. One hour before imaging, culturing medium was replaced with F12 medium (Sigma-Aldrich), supplemented with FBS, penicillin/streptomycin, and glutamine. Photoactivation was performed on a Nikon A1r confocal laser scanning microscope equipped with a Å∼60 oil immersion objective and a 37 °C and 5% CO2 incubation chamber. Laser wavelengths of 488 and 561 nm were used to acquire three pictures at 5 s intervals before photoactivation and 1 frame every 2 s for 3 min after photoactivation.

Excitation of photoactivatable GFP molecules was achieved by means of a 405 nm laser (15% power), on a single ROI matching the size of an mCherry-positive focal adhesion. Using Imaris combined with MATLAB scripts, we identified the area of photoactivation, automatically repositioned it in case of lateral shift according to the mCherry signal, and extracted the mean green intensity within the ROI for each time point. The first three time points, corresponding to the background, were averaged and subtracted from the full-time course. The intensity of the first acquisition after photoactivation was set to “100% intensity” and all other values were calculated as ratio. For each photoactivation time series, a constrained double decaying exponentials model (decreasing form) with the formula 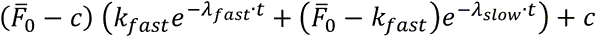 was used to fit the data points. For each of them, the half-life was defined as the time needed to lose 50% of the intensity at time 0, shown in the box and whisker plot, and used for statistical comparison. The same constrained double decaying exponentials model was also used to fit the average data, offering a visual inspection of the goodness of the fit (theoretical model). To address the heterogeneity of paxillin recruitment and stability at focal adhesions, we excluded the first 6 seconds immediately after photoactivation from our analysis. Previous studies suggest that focal adhesions form a shell-like structure where multiple low-affinity interactions cause an initial, very short-lived recruitment phase^75^. Consistent with this, we observed a quickly dissipating subpopulation of paxillin leaving the adhesion area right after activation. By excluding this early phase, we aimed to minimize the influence of this unstable pool and focus on the more stably associated population. Evidence for different paxillin subpopulations, including a membrane-associated fraction^32^, supports this approach, which may explain the fast-dissociating component.

### Spheroid assay: embedded in collagen I

Spheroids with SW480 cells were prepared following a protocol similar to the classical agarose cushion protocol^76^. We coated 96-well plates with 100 µL of a polymethyl siloxane preparation (Sylgard 184, Dow Corning), and left them overnight at 65°C to create a soft U-shape bottom well. Plates were sterilized by rinsing them with 70% w/v ethanol in deionized water and exposed to UV lamps for 1h in a biosafety cabinet (Safe 2020, Thermo Scientific). Each well was filled with 100 µL of a 5% w/v solution of Pluronic® F-127 (P2443, Sigma-Aldrich) in DPBS (14190-094, Gibco) for at least 30 minutes, to prevent subsequent cell adhesion. After discarding the Pluronic solution, each well was filled with 100 µL of culture medium containing 2000 cells. Plates were incubated at 37°C and 5% CO_2_ for 4 days. Spheroids were individually harvested using wide bore pipette tips (Axygen), deposed on 35 mm cell-culture dish on ice, and embedded in a drop of 1.5mg/mL rat tail type I collagen solution (#5153, Advanced Biomatrix) in DPBS. A 3-5 µL droplet of collagen solution containing the spheroid was then pipetted at the center of a glass bottom well of a 96-well plate (P96-1.5h-N, Cellvis) kept on ice. Once the whole plate was loaded with dome-like droplets containing spheroids, it was inverted and kept 9 minutes in the incubator at 37°C. Then the plate was put back upright at room temperature (RT) and each well filled with RT growth medium containing 500 ng ml−1 doxycycline to activate transgene expression. This method ensured spheroids were fully embedded at the center of a collagen droplet, away from the bottom of the well. Spheroids were cultured at 37°C and 5% CO_2_ for 4-5 days.

### Spheroid imaging and analysis

Each spheroid was imaged at day 4 (shROCK experiment) or at day 5 (POU5F1B mutants’ experiment) using a Leica SP8 inverted confocal microscope equipped with a motorized stage and turret, through a 25x objective (HC FLUOTAR L 25x/0,95 W VISIR, numerical aperture 0.95, water immersion). Excitation light was emitted by a Supercontinuum White Light Laser and composed of 480nm wavelength and 670 nm wavelength. Reflected and fluorescent signals were collected by the objective, filtered into two bandwidths (483 to 600nm for GFP signal, and 664 to 674 for the reflected signal), each captured by distinct photomultipliers. A third photomultiplier captured the transmitted light to obtain an image with the light traversing the samples. Each spheroid was scanned with a spatial resolution dx = dy = 0.192 µm and dz = 5µm, with a voxel dwell time of 125 ns. All image acquisition was performed using the software LAS X (Leica) and images exported in the native proprietary file. Images were analyzed using ImageJ (v 1.54). For each stack, the contrast was enhanced (saturation of 0.35% of the pixels over the whole stack) and the contour of the spheroid was segmented manually on the transmitted light images, after selecting manually the slice corresponding to the median plane of the spheroid. The segmentation was repeated similarly for the maximum extent of the cell evasion. Fluorescent and reflected light images were used to identify the most relevant frame in ambiguous cases. Spheroids that were in contact with the bottom of the wells were excluded from the analysis.

### Spheroid assay: in low or high viscosity medium

Spheroids with SW480 cells were generated out of one single cell as previously described^34^. Briefly, growth-factor-reduced Matrigel (Corning) was diluted 1:3 with cold L15 medium supplemented with 10% FBS and 1% penicillin–streptomycin. 50 µl of the diluted Matrigel were transferred to each well of a 96-well plate and allowed to polymerize at 37 °C and 5% CO_2_ for 1 h. Cells were collected from culture flasks, resuspended in diluted Matrigel and gently dispensed (2,000 cells per 50 µl) on top of the previously polymerized Matrigel in each well. After incubation at 37 °C and 5% CO_2_ for 2 h, 100 µl of L15 was added into each well. The medium was replaced every 2 days until spheroids were collected for experiments 12 days later. Spheroids were collected after dissolving the Matrigel in cold L15 and centrifuged at 2,000g for 5 min. After removing the supernatant, spheroids were resuspended in either 0.77 or 8 cP medium and plated onto collagen-I-coated (20 μg ml−1) glass bottom 96-well plates (P96-1.5h-N, Cellvis). After 60 min, the spheroids were imaged every 17 min for 21 h on the PerkinElmer Operetta high throughput microscope in brightfield, with a 10x air objective. Data was acquired with the Harmony 4.9 software. First-cell dissociation times were defined by manually recording the time required for the first cell to fully detach from each intact spheroid.

### Low-throughput BVdU treatment curves

SW480 cells stably transduced with bicistronic lentiviruses encoding (i) DCK, or POU5F1B-DCK and (ii) GFP, as well as POU5F1B-expressing and wild type SW480 control cells, were seeded into 96-well plates at 2000 cells per well in 100 μL of media. The next day, 1 M BVdU dissolved in DMSO was diluted into media to prepare 6× stock solutions of BVdU at concentrations of 640 mM, 320 mM, 160 mM, 80 mM, 40 mM and 4 mM. For each stock solution, DMSO concentration was adjusted to a final concentration of 0.25%. 0.3 μL/well of the 6× stock solutions were dispensed in the 6 first rows of the 96-well dish to achieve final concentrations of 1600 μM, 800 μM, 400 μM, 200 μM, 100 μM and 10 μM respectively. Media with 0.25% DMSO was added to the seventh row as a control. Four days later, cell viability was assessed with PrestoBlue™ reagent (A13262, Invitrogen) following manufacturer’s instructions.

### High-throughput screening using the DCK*-based positive selection assay

High-throughput chemical library screening were performed as previously described^41^. Briefly, POU5F1B-DCK SW480 cells were seeded into 384-well plates (Corning, #3764) at a density of 750 cells per well in a volume of 30 μl of media. Plates already contained a library of 5,758 lyophilized known bioactive compounds, to which cells were exposed for 24 hours before adding BVdU at a final concentration of 200 uM into columns 2 to 24. After 4 days, the GFP fluorescence of each assay plates was quantified using an Acumen scanning laser cytometer. For each plate, the average and SD of the GFP fluorescence of wells in columns 3 to 22 were calculated. For each well on an assay plate, the GFP fluorescence was converted to a z score using the formula: z(well) = [GFP (well) − μ GFP (plate)] / σ GFP (plate), where μ GFP (plate) is the mean GFP fluorescence for that plate and σ GFP (plate) is the SD for that plate.

### Radiolabeling 35S-cys/met incorporation

For metabolic labeling, transduced cells were washed with methionine /cysteine free medium, incubated 20min pulse at 37°C with 50 mCi/ml 35S-methionine/cysteine, washed and further incubated for different times at 37°C in complete medium with a 10-fold excess of non-radioactive methionine and cysteine. Proteins were immunoprecipitated and analyzed by SDS-PAGE. Autoradiography and western blot were quantified using the Typhoon Imager (Image QuantTool, GE healthcare).

---

**Supplementary Figure 1.**
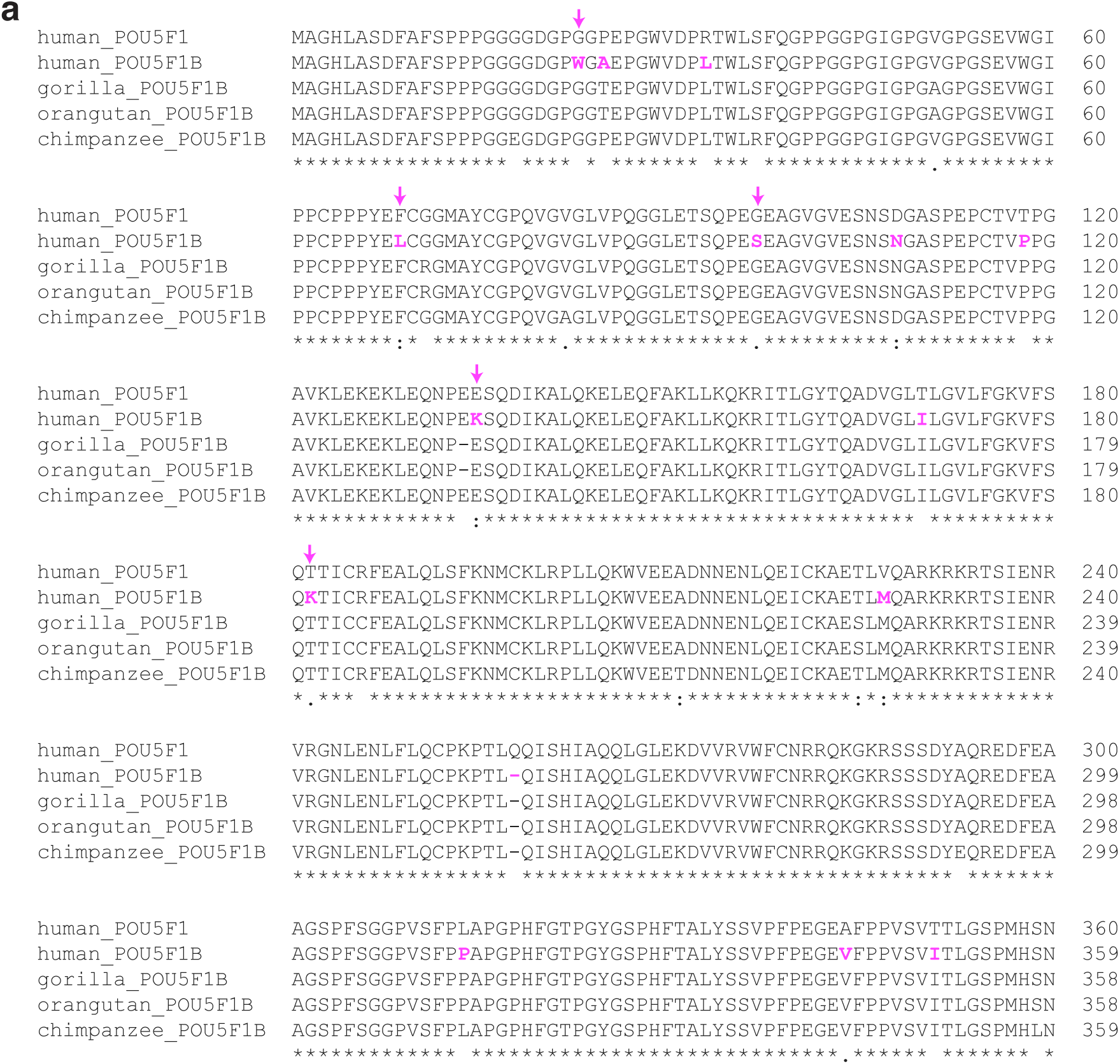
Comparing POU5F1 and POU5F1B. **a,** Amino acid sequence alignment of human POU5F1 (360 aa), human POU5F1B (359 aa), and POU5F1B from gorilla, organgutan and chimpanzee POU5F1B (358, 358 and 359 aa respectively). POU5F1/OCT4 and POU5F1B differ by 15 residues (in pink), 5 of which are unique to human POU5F1B (pink arrows).

**Supplementary Figure 2.**
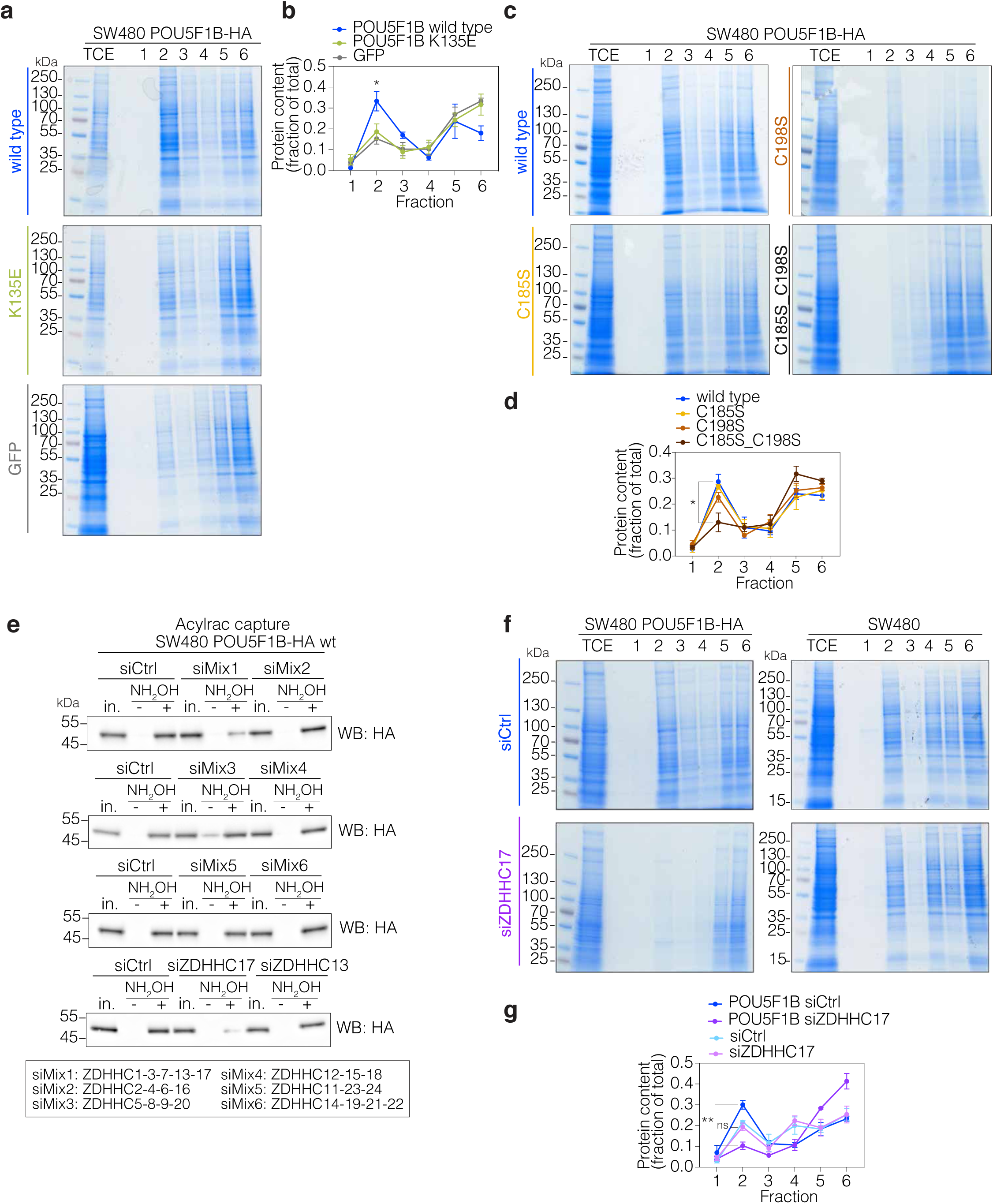
ZDHHC17-mediated S-acylation is necessary for POU5F1B to increase the protein content of membrane nanodomains. **a,** Isolation of detergent-resistant membranes (DRMs) from POU5F1B-HA-wild-type-, -K135E-, and GFP-overexpressing SW480 cells, fractions were run on SDS-PAGE and gels were stained with Coomassie. 1-3 correspond to insoluble fractions, 4-6 to soluble fractions, DRMs being traditionally found in fraction 2. **b,** Quantification of coomassie staining in (a) (Fraction 2 wild type vs. K135E P=0.04; vs.GFP P=0.033 by t-test). **c,** Same as in (a) from POU5F1B-HA-wild-type-, -C185S-, -C198S-, and -C185S_C198S-overexpressing SW480 cells. **d,** Quantification of (c) (Fraction 2 in wild type vs. C185S P=0.561; vs.C198S P=0.163; C185S_C198S P=0.029 by t-test). **e,** Acylrac capture assay in siCtrl-, siRNA mix 1 to 6-, siZDHHC17-, and siZDHHC13-POU5F1B-HA SW480 cells. **f,** Same as in (a and c) from siCtrl- and siZDHHC17-POU5F1B-HA SW480 cells and SW480 wild type cells. **g,** Quantification of (f) (Fraction 2: POU5F1B siCtrl vs. POU5F1B siZDHHC17 P=0.002; siCtrl vs. siZDHHC17 P=ns by t-test). ns not significant, * pvalue<0.05, ** pvalue<0.01.

**Supplementary Figure 3.**
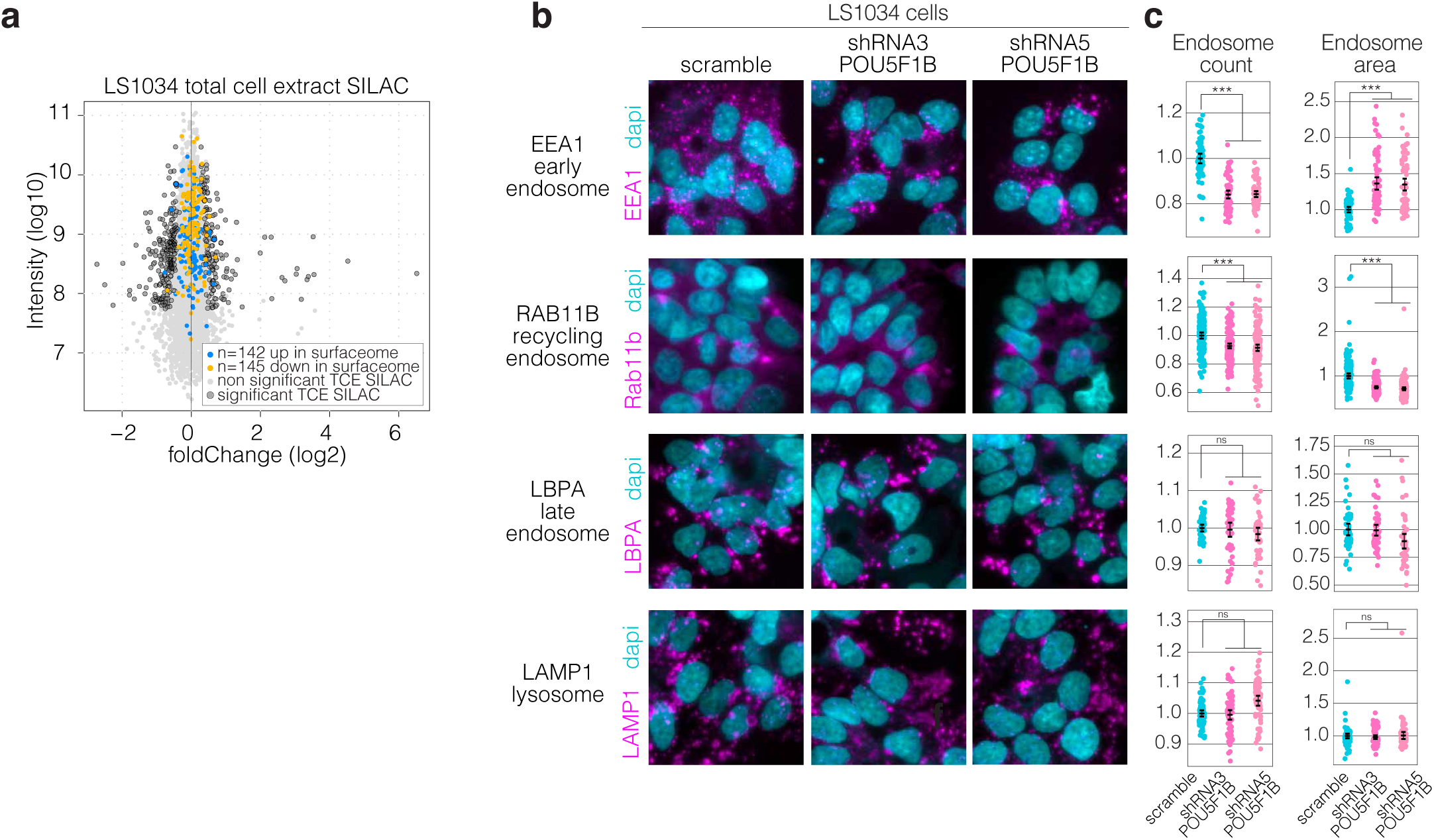
POU5F1B triggers a redistribution of proteins towards the cell surface and stimulates the early-recycling endosomal pathway. **a,** MA plot depicting relative abundance and average intensity of individual proteins identified by SILAC in total cell extracts (TCE) of sh-scramble vs. shRNA3 & shRNA5 LS1034 cells (data from https://doi.org/10.1038/s41467-022-32649-7). Measurements were performed in independent duplicates, each dot represents a detected protein, with significantly changed ones (P < 0.05) circled in black. 142 upregulated and 145 downreguated proteins in the surfaceome from the same cells (Fig 3F) are colored in blue and yelow respectively. **b,** Representative immunofluorescence-confocal microscopy of sh-scramble, shRNA3, and shRNA5 LS1034 cells, with EEA1, RAB11B, LBPA, and LAMP1 lysosomes in magenta and the nuclear marker dapi in cyan. **c,** Endosome count and area quantification from cells in (b) (Respective count and area from EEA1 P=1.34e-33 P=3.27e-13; RAB11B P=6.94e-10 P=3.87e-25; LBPA P=0.29 P=0.11; LAMP1 P=0.06 P=0.82 positive endosomes in scramble vs. sh3&sh5 cells). ns, not significant, *** pvalue<0.001

**Supplementary Figure 4.**
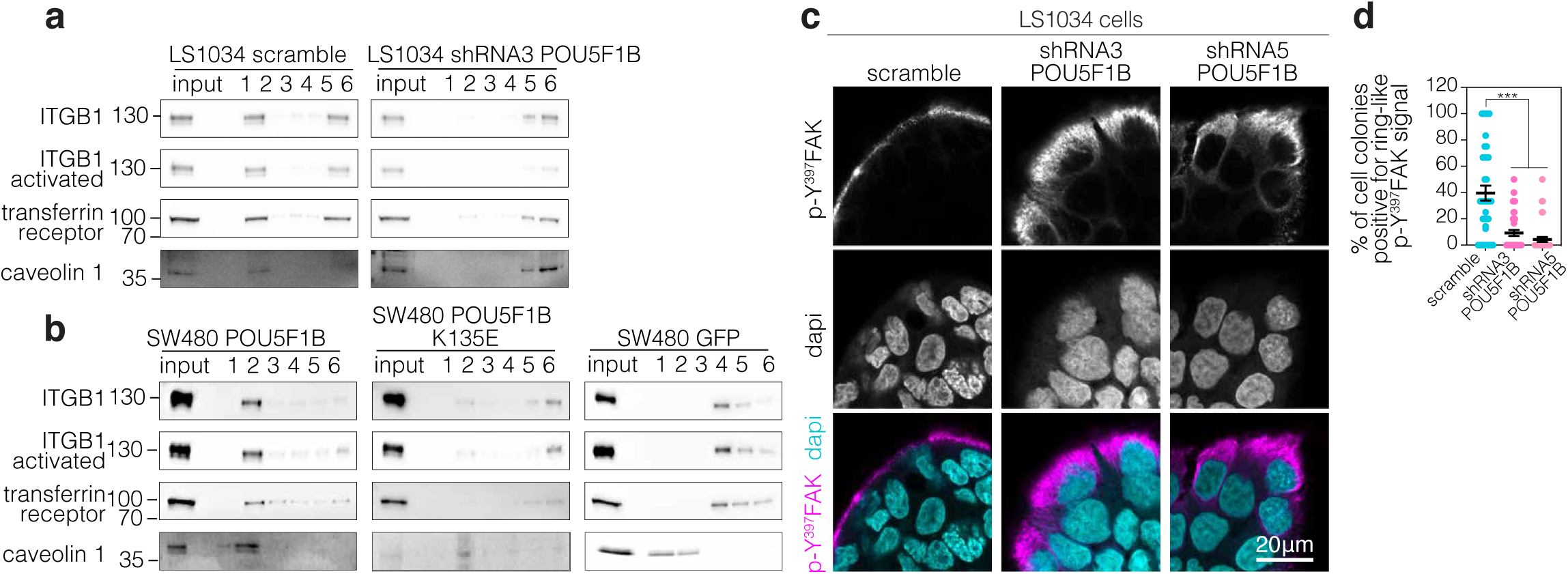
POU5F1B induces accumulation of ITGB1 in DRMs and activated focal adhesion kinase (p-Y^397^ FAK) at the periphery of colorectal cancer cells. **a,** Western blot analysis of ITGB1 and activated ITGB1 in detergent-resistant membranes (DRMs) extraction from sh-scramble, shRNA3 & shRNA5 LS1034 cells, and **b,** POU5F1B-, POU5F1B-K135E- and GFP-overexpressing SW480 cells. Caveolin1 and transferrin receptor are used as controls for insoluble and soluble fractions, respectively. **c,** Representative immunofluorescence-confocal microscopy of cells presented in (a), with p-Y^397^ FAK in magenta and the nuclear marker dapi in cyan. **d,** Quantification of the percentage of cell colonies positive for peripheral ring-like p-Y^397^FAK signal from cells in (c) (scramble vs. shRNA3 P=1.67e-05; vs. shRNA5 P=5.0e-08 by t-test). *** pvalue<0.001

**Supplementary Figure 5.**
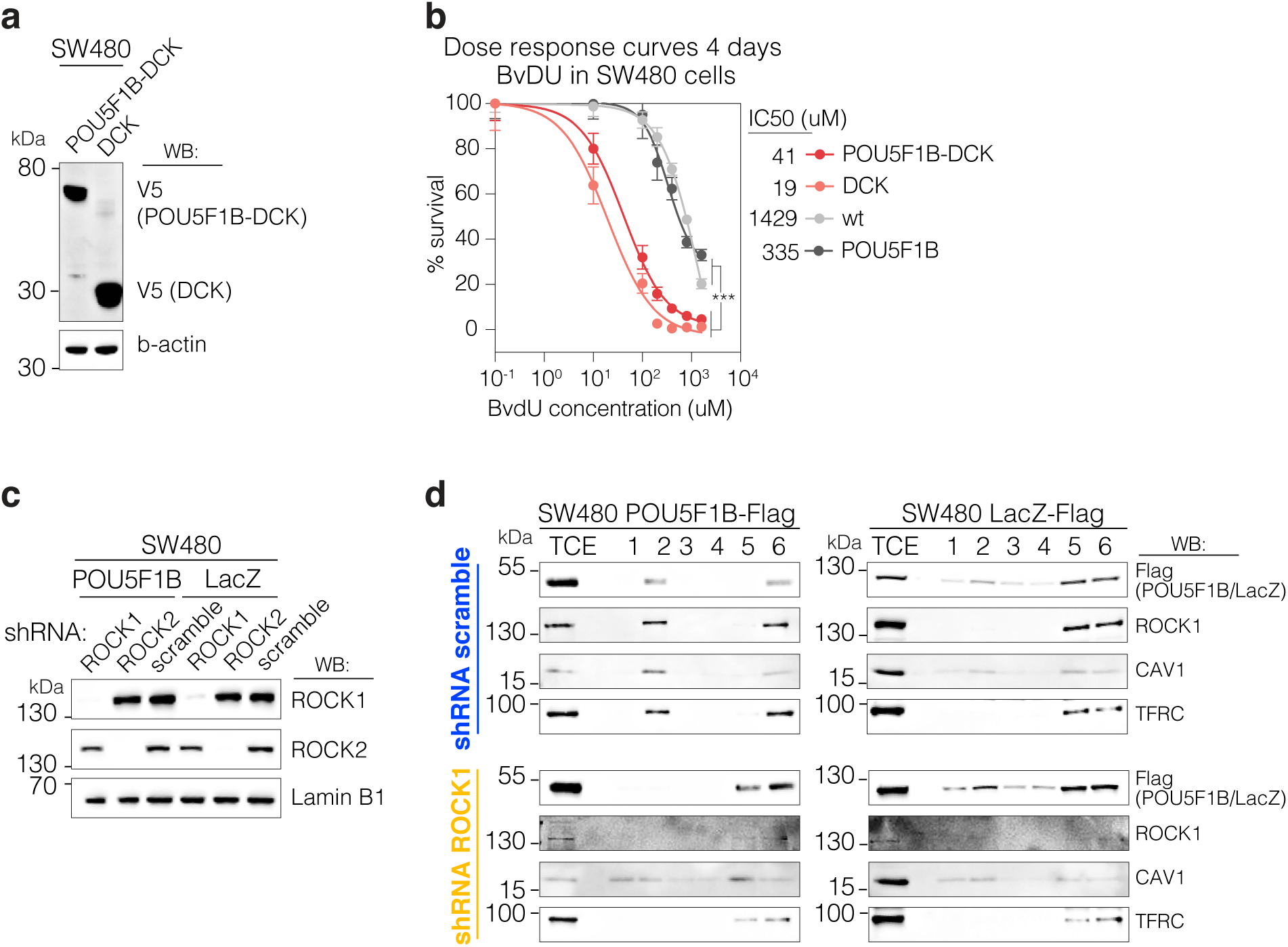
Screening for inducers of POU5F1B degradation and examining their impact on DRMs. **a,** Immunoblot analysis of SW480 cells infected with the lentiviral vectors depicted in (Fig. 6A). **b,** 2-bromovinyldeoxyuridine (BvdU) dose response curves in wild type, DCK-, POU5F1B-DCK-, and POU5F1B-expressing SW480 cells, 4 days post-treatment (P = 9.7e-27 by extra sum-of-squares F-test). **c,** Immunoblot analysis of POU5F1B- and LacZ-expressing SW480 cells infected with lentiviral vectors expressing doxycycline-inducible shRNAs against ROCK1, ROCK2 and scramble sequences. **d,** Immunoblot analysis of ROCK1 and Flag-tagged POU5F1B and LacZ in detergent-resistant membranes (DRMs) extraction from sh-scramble and shROCK1 SW480 cells expressing either POU5F1B or LacZ. Caveolin1 and transferrin receptor are used as controls for insoluble and soluble fractions, respectively. *** pvalue<0.001.

