## supplementary_figure_1 for "The human-restricted oncoprotein POU5F1B enhances cell invasiveness through plasma membrane remodeling"

**a**

```
human_POU5F1         MAGHLASDFAFSPPPGGGGDGPGGPEPGWVDPRTWLSFQGPPGGPGIGPGVGPGSEVWGI  60
human_POU5F1B        MAGHLASDFAFSPPPGGGGDGPWGAEPGWVDPLTWLSFQGPPGGPGIGPGVGPGSEVWGI  60
gorilla_POU5F1B      MAGHLASDFAFSPPPGGGGDGPGGTEPGWVDPLTWLSFQGPPGGPGIGPGAGPGSEVWGI  60
orangutan_POU5F1B    MAGHLASDFAFSPPPGGGGDGPGGTEPGWVDPLTWLSFQGPPGGPGIGPGAGPGSEVWGI  60
chimpanzee_POU5F1B   MAGHLASDFAFSPPPGGEGDGPGGPEPGWVDPLTWLRFQGPPGGPGIGPGVGPGSEVWGI  60
                     ***************** **** * ******* *** *************.*********

human_POU5F1         PPCPPPYEFCGGMAYCGPQVGVGLVPQGGLETSQPEGEAGVGVESNSDGASPEPCTVTPG  120
human_POU5F1B        PPCPPPYELCGGMAYCGPQVGVGLVPQGGLETSQPESEAGVGVESNSNGASPEPCTVPPG  120
gorilla_POU5F1B      PPCPPPYEFCRGMAYCGPQVGVGLVPQGGLETSQPEGEAGVGVESNSNGASPEPCTVPPG  120
orangutan_POU5F1B    PPCPPPYEFCRGMAYCGPQVGVGLVPQGGLETSQPEGEAGVGVESNSNGASPEPCTVPPG  120
chimpanzee_POU5F1B   PPCPPPYEFCGGMAYCGPQVGAGLVPQGGLETSQPEGEAGVGVESNSDGASPEPCTVPPG  120
                     ********:* **********.**************.**********:********* **

human_POU5F1         AVKLEKEKLEQNPEESQDIKALQKELEQFAKLLKQKRITLGYTQADVGLTLGVLFGKVFS  180
human_POU5F1B        AVKLEKEKLEQNPEKSQDIKALQKELEQFAKLLKQKRITLGYTQADVGLILGVLFGKVFS  180
gorilla_POU5F1B      AVKLEKEKLEQNP-ESQDIKALQKELEQFAKLLKQKRITLGYTQADVGLILGVLFGKVFS  179
orangutan_POU5F1B    AVKLEKEKLEQNP-ESQDIKALQKELEQFAKLLKQKRITLGYTQADVGLILGVLFGKVFS  179
chimpanzee_POU5F1B   AVKLEKEKLEQNPEESQDIKALQKELEQFAKLLKQKRITLGYTQADVGLILGVLFGKVFS  180
                     ************* :********************************** **********

human_POU5F1         QTTICRFEALQLSFKNMCKLRPLLQKWVEEADNNENLQEICKAETLVQARKRKRTSIENR  240
human_POU5F1B        QKTICRFEALQLSFKNMCKLRPLLQKWVEEADNNENLQEICKAETLMQARKRKRTSIENR  240
gorilla_POU5F1B      QTTICCFEALQLSFKNMCKLRPLLQKWVEEADNNENLQEICKAESLMQARKRKRTSIENR  239
orangutan_POU5F1B    QTTICCFEALQLSFKNMCKLRPLLQKWVEEADNNENLQEICKAESLMQARKRKRTSIENR  239
chimpanzee_POU5F1B   QTTICRFEALQLSFKNMCKLRPLLQKWVEETDNNENLQEICKAETLMQARKRKRTSIENR  240
                     *.*** ************************:*************:*:*************

human_POU5F1         VRGNLENLFLQCPKPTLQQISHIAQQLGLEKDVVRVWFCNRRQKGKRSSSDYAQREDFEA  300
human_POU5F1B        VRGNLENLFLQCPKPTL-QISHIAQQLGLEKDVVRVWFCNRRQKGKRSSSDYAQREDFEA  299
gorilla_POU5F1B      VRGNLENLFLQCPKPTL-QISHIAQQLGLEKDVVRVWFCNRRQKGKRSSSDYAQREDFEA  298
orangutan_POU5F1B    VRGNLENLFLQCPKPTL-QISHIAQQLGLEKDVVRVWFCNRRQKGKRSSSDYAQREDFEA  298
chimpanzee_POU5F1B   VRGNLENLFLQCPKPTL-QISHIAQQLGLEKDVVRVWFCNRRQKGKRSSSDYEQREDFEA  299
                     ***************** ********************************** *******

human_POU5F1         AGSPFSGGPVSFPLAPGPHFGTPGYGSPHFTALYSSVPFPEGEAFPPVSVTTLGSPMHSN  360
human_POU5F1B        AGSPFSGGPVSFPPAPGPHFGTPGYGSPHFTALYSSVPFPEGEVFPPVSVITLGSPMHSN  359
gorilla_POU5F1B      AGSPFSGGPVSFPPAPGPHFGTPGYGSPHFTALYSSVPFPEGEVFPPVSVITLGSPMHSN  358
orangutan_POU5F1B    AGSPFSGGPVSFPPAPGPHFGTPGYGSPHFTALYSSVPFPEGEVFPPVSVITLGSPMHSN  358
chimpanzee_POU5F1B   AGSPFSGGPVSFPLAPGPHFGTPGYGSPHFTALYSSVPFPEGEVFPPVSVITLGSPMHLN  359
                     ************* *****************************.****** ******* *
```

**Supplementary Figure 1. Comparing POU5F1 and POU5F1B. a,** Amino acid sequence alignment of human POU5F1 (360 aa), human POU5F1B (359 aa), and POU5F1B from gorilla, organgutan and chimpanzee POU5F1B (358, 358 and 359 aa respectively). POU5F1/OCT4 and POU5F1B differ by 15 residues (in pink), 5 of which are unique to human POU5F1B (pink arrows).
