## supplementary_figure_2 for "The human-restricted oncoprotein POU5F1B enhances cell invasiveness through plasma membrane remodeling"

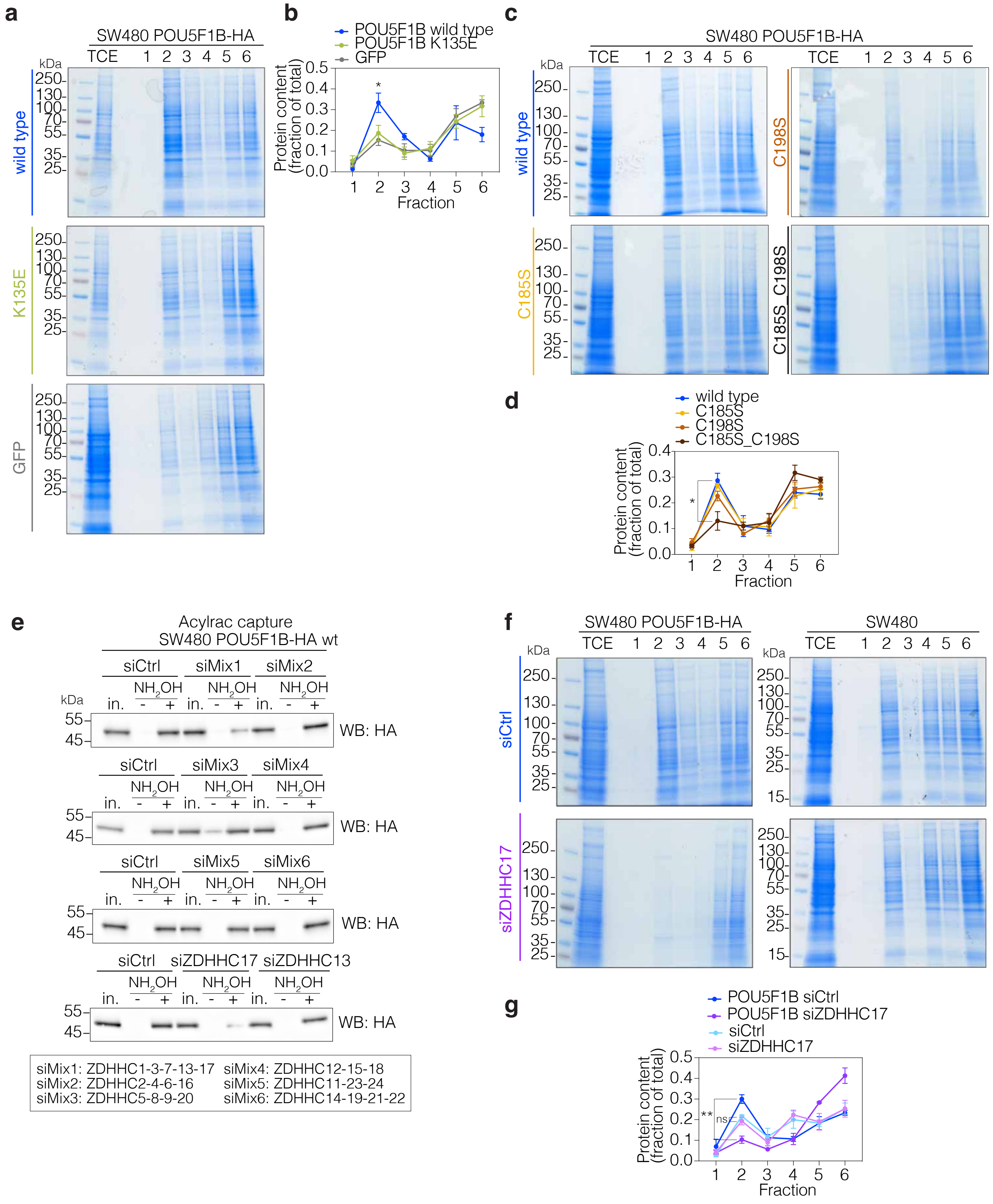


**Supplementary Figure 2.ZDHHC17-mediated S-acylation is necessary for POU5F1B to increase the protein content of membrane nanodomains. a,** Isolation of detergent-resistant membranes (DRMs) from POU5F1B-HA-wild-type-, -K135E-, and GFP-overexpressing SW480 cells, fractions were run on SDS-PAGE and gels were stained with Coomassie. 1-3 correspond to insoluble fractions, 4-6 to soluble fractions, DRMs being traditionally found in fraction 2. **b,** Quantification of coomassie staining in (a) (Fraction 2 wild type vs. K135E P=0.04; vs.GFP P=0.033 by t-test). **c,** Same as in (a) from POU5F1B-HA-wild-type-, -C185S-, -C198S-, and -C185S_C198S-overexpressing SW480 cells. **d,** Quantification of (c) (Fraction 2 in wild type vs. C185S P=0.561; vs.C198S P=0.163 ; C185S_C198S P=0.029 by t-test). **e,** Acylrac capture assay in siCtrl-, siRNA mix 1 to 6-, siZDHHC17-, and siZDHHC13-POU5F1B-HA SW480 cells. **f,** Same as in (a and c) from siCtrl- and siZDHHC17- POU5F1B-HA SW480 cells and SW480 wild type cells. **g,** Quantification of (f) (Fraction 2: POU5F1B siCtrl vs. POU5F1B siZDHHC17 P=0.002; siCtrl vs. siZDHHC17 P=ns by t-test). ns not significant, * pvalue<0.05, ** pvalue<0.01.
