## supplementary_figure_3 for "The human-restricted oncoprotein POU5F1B enhances cell invasiveness through plasma membrane remodeling"

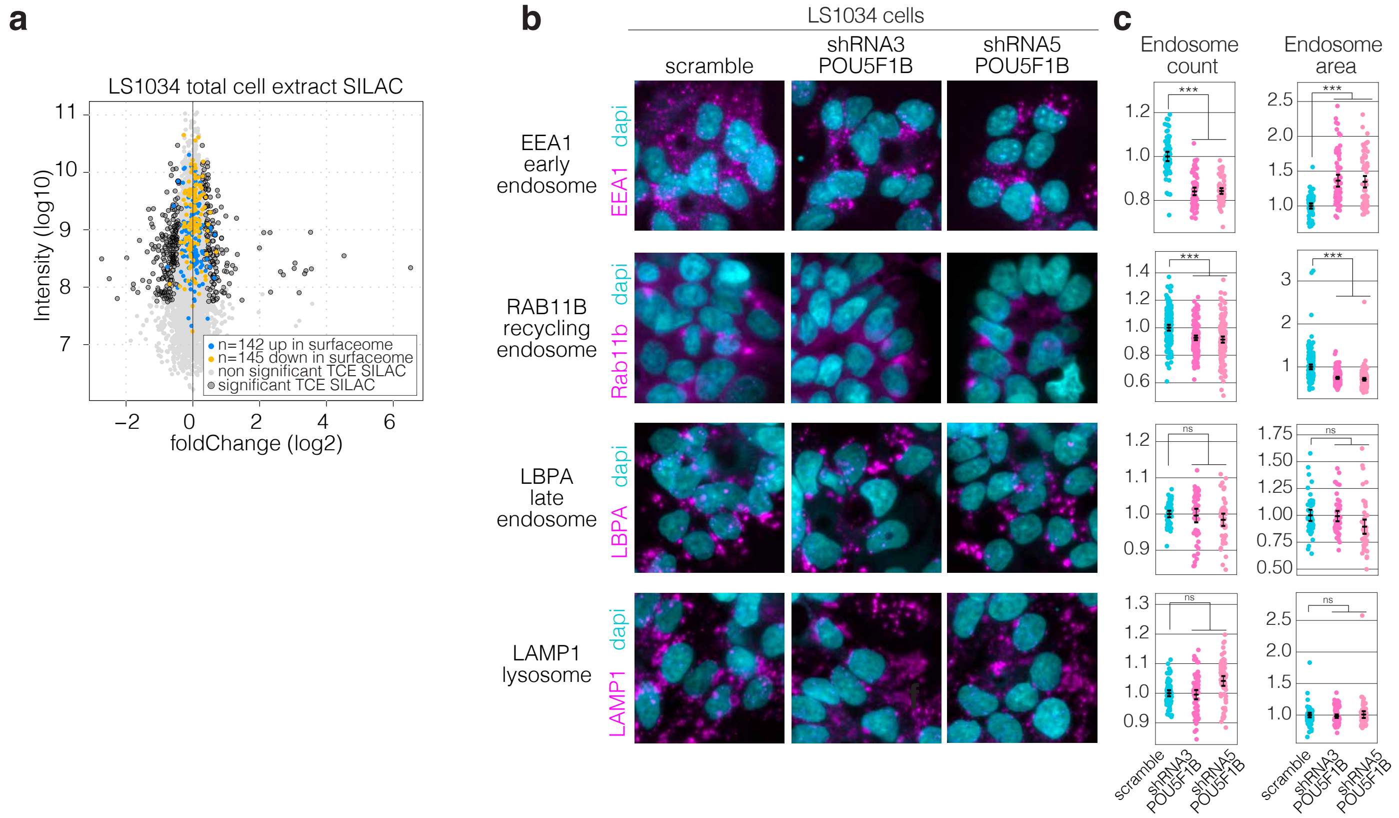


**Supplementary Figure 3. POU5F1B triggers a redistribution of proteins towards the cell surface and stimulates the early-recycling endosomal pathway. a,** MA plot depicting relative abundance and average intensity of individual proteins identified by SILAC in total cell extracts (TCE) of sh-scramble vs. shRNA3 & shRNA5 LS1034 cells (data from https://doi.org/10.1038/s41467-022-32649-7). Measurements were performed in independent duplicates, each dot represents a detected protein, with significantly changed ones (P < 0.05) circled in black. 142 upregulated and 145 downreguated proteins in the surfaceome from the same cells (Fig 3F) are colored in blue and yelow respectively. **b,** Representative immunofluorescence-confocal microscopy of sh-scramble, shRNA3, and shRNA5 LS1034 cells, with EEA1, RAB11B, LBPA, and LAMP1 lysosomes in magenta and the nuclear marker dapi in cyan. **c,** Endosome count and area quantification from cells in (b) (Respective count and area from EEA1 P=1.34e-33 P=3.27e-13; RAB11B P=6.94e-10 P=3.87e-25; LBPA P=0.29 P=0.11; LAMP1 P=0.06 P=0.82 positive endosomes in scramble vs. sh3&sh5 cells). ns, not significant, *** pvalue<0.001
