## supplementary_figure_4 for "The human-restricted oncoprotein POU5F1B enhances cell invasiveness through plasma membrane remodeling"

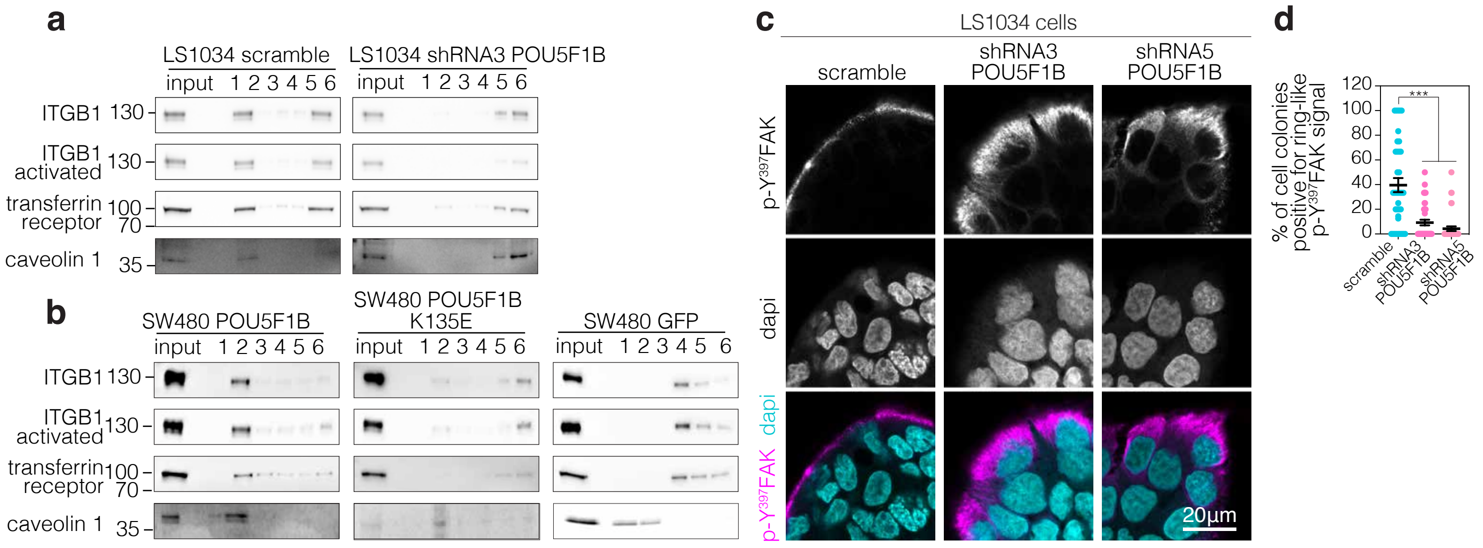


**Supplementary Figure 4. POU5F1B induces accumulation of ITGB1 in DRMs and activated focal adhesion kinase (p-Y$^{397}$ FAK) at the periphery of colorectal cancer cells. a,** Western blot analysis of ITGB1 and activated ITGB1 in detergent-resistant membranes (DRMs) extraction from sh-scramble, shRNA3 & shRNA5 LS1034 cells, and **b,** POU5F1B-, POU5F1B-K135E- and GFP-overexpressing SW480 cells. Caveolin1 and transferrin receptor are used as controls for insoluble and soluble fractions, respectively. **c,** Representative immunofluorescence-confocal microscopy of cells presented in (a), with p-Y$^{397}$ FAK in magenta and the nuclear marker dapi in cyan. **d,** Quantification of the percentage of cell colonies positive for peripheral ring-like p-Y$^{397}$FAK signal from cells in (c) (scramble vs. shRNA3 P=1.67e-05; vs. shRNA5 P=5.0e-08 by t-test). *** pvalue<0.001
