## supplementary_figure_5 for "The human-restricted oncoprotein POU5F1B enhances cell invasiveness through plasma membrane remodeling"

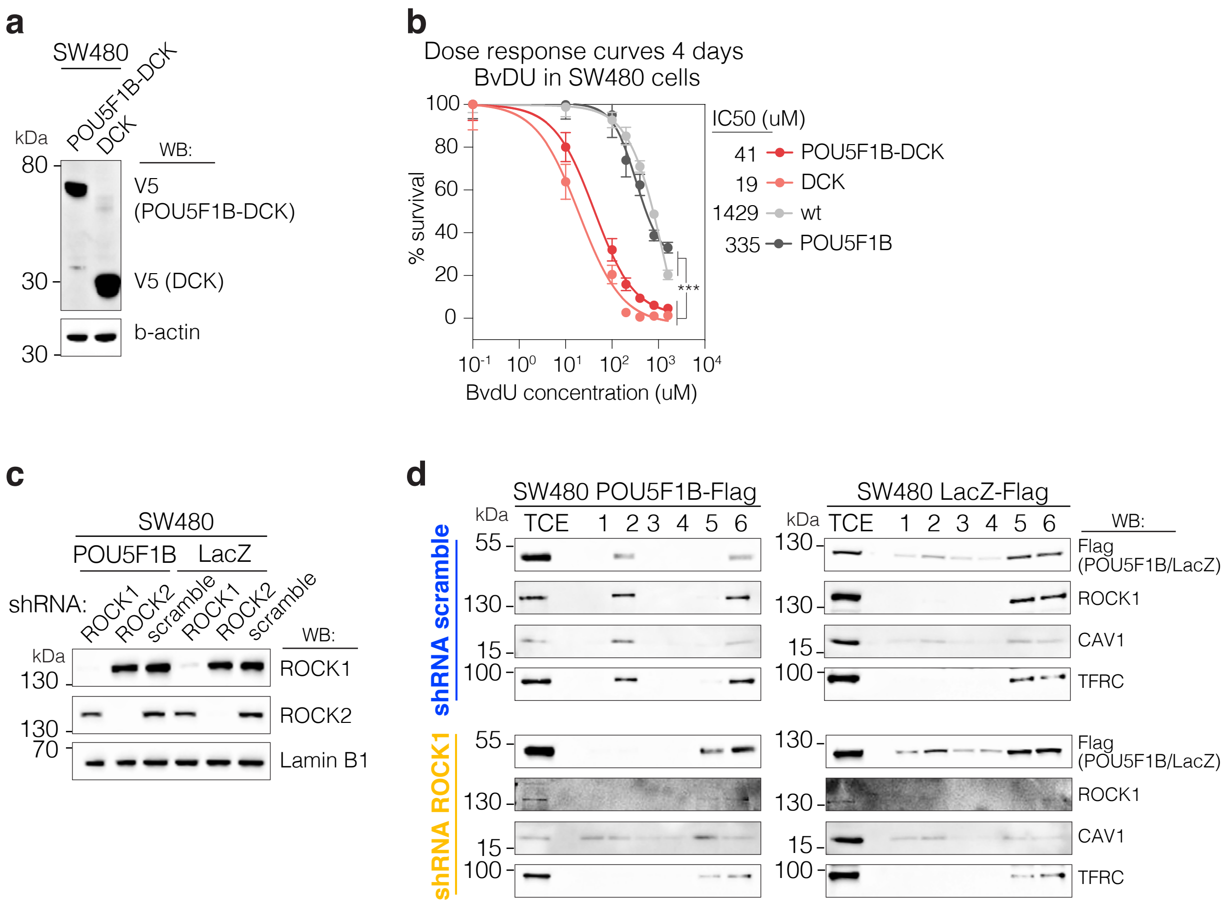


**Supplementary Figure 5. Screening for inducers of POU5F1B degradation and examining their impact on DRMs. a,** Immunoblot analysis of SW480 cells infected with the lentiviral vectors depicted in (Fig. 6A). **b,** 2-bromovinyldeoxyuridine (BvdU) dose response curves in wild type, DCK-, POU5F1B-DCK-, and POU5F1B-expressing SW480 cells, 4 days post-treatment (P = 9.7e-27 by extra sum-of-squares F-test). **c,** Immunoblot analysis of POU5F1B- and LacZ-expressing SW480 cells infected with lentiviral vectors expressing doxycycline-inducible shRNAs against ROCK1, ROCK2 and scramble sequences. **d,** Immunoblot analysis of ROCK1 and Flag-tagged POU5F1B and LacZ in detergent-resistant membranes (DRMs) extraction from sh-scramble and shROCK1 SW480 cells expressing either POU5F1B or LacZ. Caveolin1 and transferrin receptor are used as controls for insoluble and soluble fractions, respectively. *** pvalue<0.001.
